# Shape-restrained modelling of protein-small molecule complexes with HADDOCK

**DOI:** 10.1101/2021.06.10.447890

**Authors:** Panagiotis I. Koukos, Manon Réau, Alexandre M. J. J. Bonvin

## Abstract

Small molecule docking remains one of the most valuable computational techniques for the structure prediction of protein-small molecule complexes. It allows us to study the interactions between compounds and the protein receptors they target at atomic detail, in a timely and efficient manner. Here we present a new protocol in HADDOCK, our integrative modelling platform, which incorporates homology information for both receptor and compounds. It makes use of HADDOCK’s unique ability to integrate information in the simulation to drive it toward conformations which agree with the provided data. The focal point is the use of shape restraints derived from homologous compounds bound to the target receptors. We have developed two protocols: In the first, the shape is composed of fake atom beads based on the position of the heavy atoms of the homologous template compound, whereas in the second the shape is additionally annotated with pharmacophore data, for some or all beads. For both protocols, ambiguous distance restraints are subsequently defined between those beads and the heavy atoms of the ligand to be docked. We have benchmarked the performance of these protocols with a fully unbound version of the widely used DUD-E dataset. In this unbound docking scenario, our template/shape-based docking protocol reaches an overall success rate of 81% on 99 complexes, which is close to the best results reported for bound docking on the DUD-E dataset.

**Table of contents graphic:** 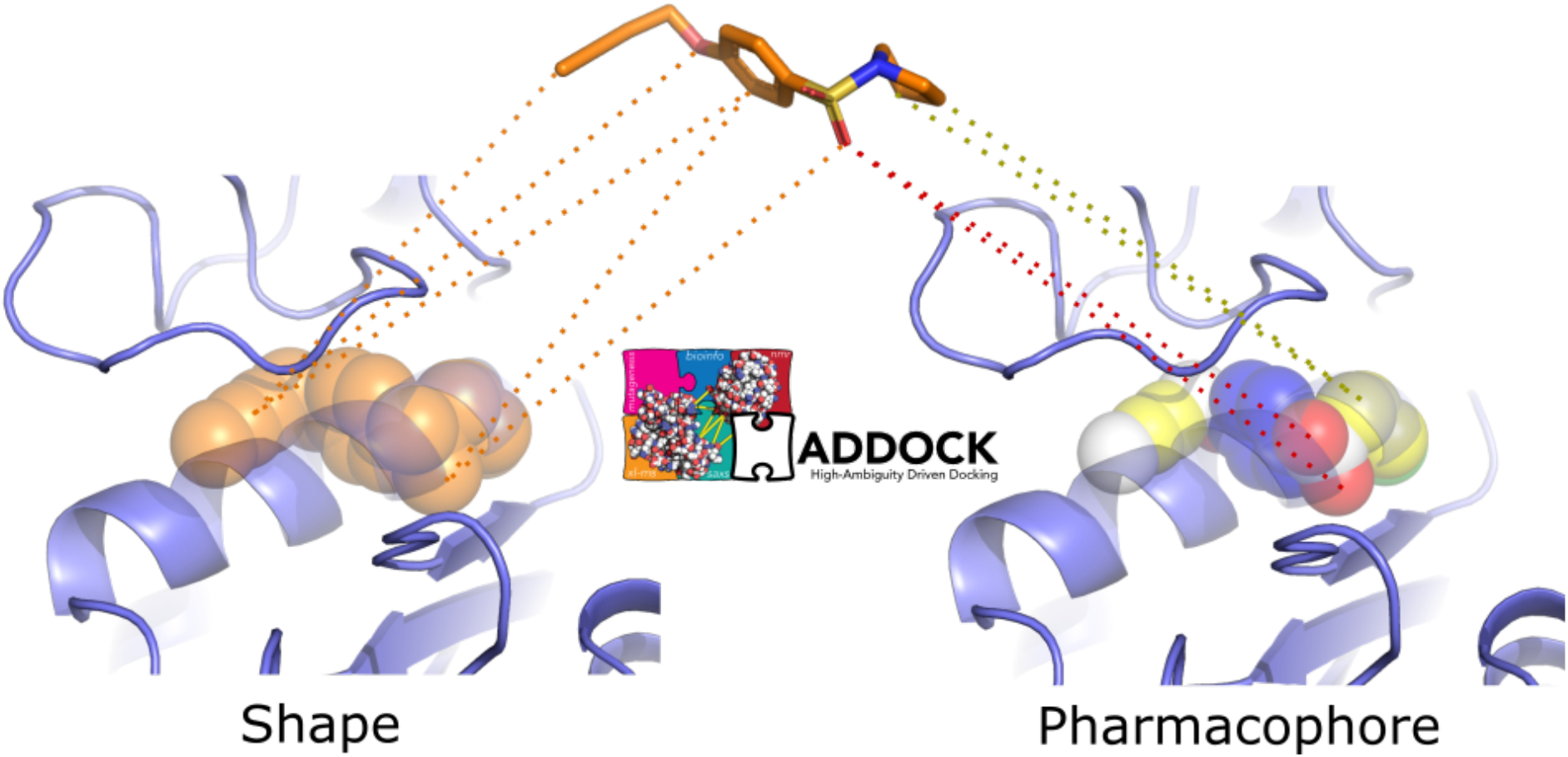

## Introduction

The importance of reliable methods for the docking of small molecule compounds to receptors of pharmaceutical interest cannot be understated. The nature of modern drug development practices dictates the gradual filtering of millions (perhaps even hundreds of millions) of compounds contained in virtual libraries to, ultimately, a few dozen lead compounds that can be further optimised before their clinical potential is investigated in animal and human trials [1,2]. This set of practices – collectively known as Computer-Aided Drug Design (CADD) – encompasses a variety of methods such as virtual screening of compounds, molecular docking with recent developments making use of machine learning-based approaches [3,4] and binding affinity prediction.

A relatively recent development in this space has been the advent of blind experiments (or exercises) focusing on pose prediction and binding affinity ranking/prediction. Examples of such initiatives are the – aptly-named – ‘Grand Challenges’ (GC), held by the Drug Design Data Resource (D3R) consortium, on an annual basis between 2015 and 2018 [5–8]. These experiments built on the experience of the earlier Community Structure-Activity Resource (CSAR) experiment [9]. The stated drives behind these initiatives have been the desire to establish – and push – state-of-the-art performance in these challenging modelling problems, develop commonly agreed-upon standards of evaluating and assessing aforementioned performance, codify best practices and facilitate communication and sharing of data between pharmaceutical companies and academic investigators. In these targets, these initiatives have been, largely, successful.

It is through our participation [10–12] with HADDOCK (**H**igh **A**mbiguity **D**riven **DOCK**ing), our integrative modelling platform [13,14], in the D3R experiment that we initially developed docking protocols tailored to the peculiarities of protein-small molecule complex modelling.

Whereas our protocol of choice for the 2016 iteration of the GC did not earn us a spot among the top performing groups of that round, it did allow us to better understand the problems specific to protein-small molecule docking. The main takeaway point was that making use of the most closely related receptor for every target compound significantly improved the outcome of the modelling. In this case, the best receptor was identified after comparing crystallographic compounds with the target compound and selecting the receptor conformation with the most similar ligand. With that knowledge we optimised additional aspects of our approach for GC 2017 – always prioritising the use of high-quality experimental information for every step of the process. That revised protocol resulted in HADDOCK submitting one of the most accurate predictions for the pose prediction component of the challenge. We applied the same protocol in GC 2018 with equally good results.

Our successful D3R protocol can be summarised as: 1) After identifying highly homologous receptors with a co-crystallised compound, we compare the similarity of all crystallographic compounds to all target compounds and select the receptor conformation whose compound has the highest similarity to the compound to dock. 2) Prior to docking, we filter the generated conformers by comparing their 3D shape with that of the most similar crystallographic compound and select the 10 closest conformers in terms of shape similarity. 3) Finally, these 10 conformers are placed into the binding pocket by superimposing their shape onto the shape of the crystallographic compound and the model was refined in HADDOCK (i.e., no docking is performed).

While the use of shapes in protein-ligand modelling is not novel, even dating back to the earliest days of the field [15,16], with common approaches emphasising shape complementarity or overlap [17–19], to the best of our knowledge, HADDOCK’s ability to drive the desired compound into specific parts of the binding pocket via the use of shape information in combination with other kinds of restraints in a unified integrative modelling framework is unique among modelling software.

Here, we present a new shape-based protocol which incorporates all the lessons that we have learned over three years of participating in the D3R blind docking experiment into a protocol designed for HADDOCK, bypassing one of the main limitations of the previous protocols – their reliance on external software for significant parts of the ligand-based modelling process. This limitation did not allow us to use the integrative modelling and flexible refinement capabilities of HADDOCK as the rigid-body and semi-flexible refinement stages were bypassed and only the final flexible refinement stage was performed. Also, the fact we are no longer dependent on commercial software for parts of the modelling workflow means we are free to develop this protocol into a pipeline accessible through the freely available HADDOCK webserver [14] (https://wenmr.science.uu.nl/haddock2.4/).

In the new shape-restrained protocols presented here, the principles underlying template identification and conformer generation procedures are similar to the ones described previously. The main difference is that, after identifying a suitable receptor template for each target, its bound compound is transformed into a shape consisting of fake beads. In the pharmacophore-based version of the protocol, a pharmacophore shape is only introduced if features can be assigned for that part of the template compound, otherwise a regular shape bead is used. Ambiguous distance restraints to guide the docking are then defined between those beads and the heavy atoms of the compound. There is no selection of conformers prior to docking, but instead, up to 50 conformers are docked into the receptor template. The most suited conformations are automatically selected during the docking by HADDOCK based on the shape restraints. These restraints are also drivers of possible conformational changes in the ligand during the flexible refinement stage of HADDOCK. We evaluate the performance of these new shape-restrained protocols on the Database of Useful Decoys-Enhanced (DUD-E) dataset that contains crystallographic structures for 102 protein-small molecule complexes [20]. Among those, 58 proteins can be classified as protein kinases, nuclear receptors, proteases, GPCRs, cleaving enzymes, cyclooxygenases, cytochromes P450, ion channels or histone deacetylases and 54 are unique representants of a protein family [20,21]. The DUD-E and its previous versions have been mostly used to benchmark virtual screening tools [22,23]. When used to benchmark sampling tools, in most cases, the bound conformations of the target proteins were used [21,24]. The use of bound structures however, biases the outcome of the docking since it suggests that we know the exact conformation of the target proteins when bound to their ligand and that we effectively ignore the intrinsic flexibility of their binding site and any binding-induced conformational changes [25]. In this work, we introduce a new unbound benchmark version of the DUD-E that allows us to assess the performance of the HADDOCK shape-restrained protocols in more realistic conditions.

## Results and Discussion

### Dataset

We successfully identified templates for 99 of the 102 targets that are part of the DUD-E dataset (Tables S1 and S2). Those templates correspond to identical or homologous receptors that display the exact same sequence at the binding site level and that are co-crystalized with a different ligand. The amount of conformational changes in the binding sites ranges from 0.08Å to 4.01Å (see Tables S1 and S2). No usable templates could be identified for targets *cxcr4*, *drd3* and *kpcb*. For *kpcb* and *drd3* we failed to identify any homologous templates whereas for cxcr4 we did identify some, but they were all in complex with the same compound as the reference receptor, so we discarded them. We identified at least one viable template for the remaining targets and selected one for docking based on the similarity of the reference and template compounds (see Methods). The similarity values range from almost 1 to below 0.2 indicating the presence of templates whose compounds are almost identical to their respective reference compounds on one end and templates that are almost entirely dissimilar on the other. The distribution of similarity values for the templates used for the two protocols can be seen in SI Figure 1. As we expected the similarity between template and reference compounds to be a limiting factor for the outcome of the modelling, we decided to investigate that relationship for the shape protocol, by identifying an additional low similarity template for all targets whose original template had a similarity value greater than 0.8 (see SI Figure 1). In this way, we can compare the performance in the subset of the dataset which includes both low- and high-similarity templates. In total, for 34 of the 35 targets for which the original similarity value was greater than 0.8 we could identify an additional, low-similarity template (no alternative templates could be identified for target *kith*).

### Conformer generation

Prior to assessing the docking performance, we evaluate the performance of the conformer generation as we expect it to have a significant impact on the outcome of the docking, since starting ligand conformations that are very different to the reference compound would need to undergo significant conformational rearrangements. In summary, RDKit and the settings we have chosen (see Methods) perform well as we are able to generate at least one acceptable pose (conformers whose heavy-atom RMSD to the reference bound ligand conformation is ≤ 2Å) for all but five target compounds. Specifically, the mean percentage of acceptable poses is 66 ± 36 (median: 78), however there are 11 targets for which less than 10% of the generated conformers are acceptable, 5 of those with no generated poses below 2Å (see SI Figure 2).

### Docking

Figure 1 shows the docking strategy behind the shape- and pharmacophore-driven protocols. In short, after selecting a suitable template, a shape based on the template compound is generated, restraints are defined between template shapes and target conformers, and docking is performed with HADDOCK (see Methods for more details).

**Figure 1:**
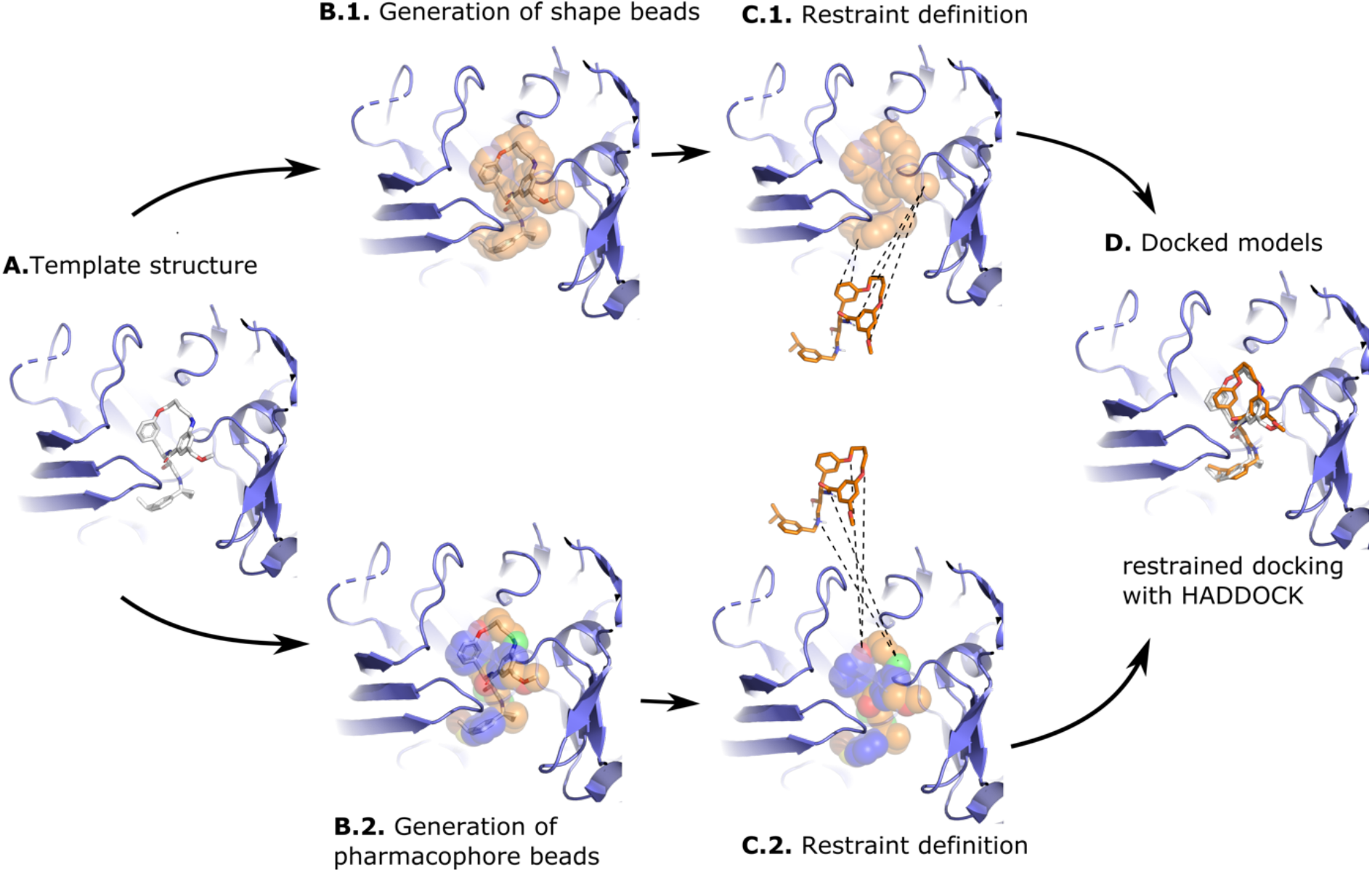
Illustration of the shape-based and the pharmacophore-based docking protocol. **Panel A** shows a suitable receptor template identified based on the similarity of its bound ligand and the ligand to be docked (see Methods). **Panel B** shows the heavy atoms of the crystallographic compound transformed into shape beads (**B.1**) or pharmacophore beads with the coloured beads representing different pharmacophore properties (**B.2**). The crystallographic compound is then removed from the pocket and restraints are defined between the beads and the conformers: In the shape-based protocol (**C.1**) restraints are defined between all atoms of the compound and all beads of the shape, while in the pharmacophore-based protocol (**C.2**), restraints are defined between atoms of the compound and beads that share identical pharmacophore features. **Panel D** shows a docked model superimposed onto the template structure. The protein receptor is shown as slate cartoon, the crystallographic compound as white sticks, the generated and docked compounds as orange sticks and the shape beads as transparent orange spheres. All molecular graphics were generated with PyMOL [26].

Figure 2 shows the success rate for the two protocols with the ‘shape’ and ‘pharm’ groups highlighting the performance of the shape- and pharmacophore-driven protocols, respectively. The success rate is defined as the percentage of targets for which a model of acceptable (or better) quality has been generated within the topN ranked models. For the shape protocol, the performance is very poor when considering only the top ranked models during the rigid body stage but improves significantly in the refinement stage increasing to 58.6% and 71.7% when considering the top1 and top5 models, respectively, with an overall success rate of 81.8% when considering all refined models. The respective values for the pharmacophore protocol are 53.5%, 63.6% and 75.8% indicating a slightly lower performance compared to the shape protocol. Note again that those success rates are for an unbound docking scenario (i.e., both ligand and receptor starting conformations deviate from their reference bound conformations in the complex).

**Figure 2:**
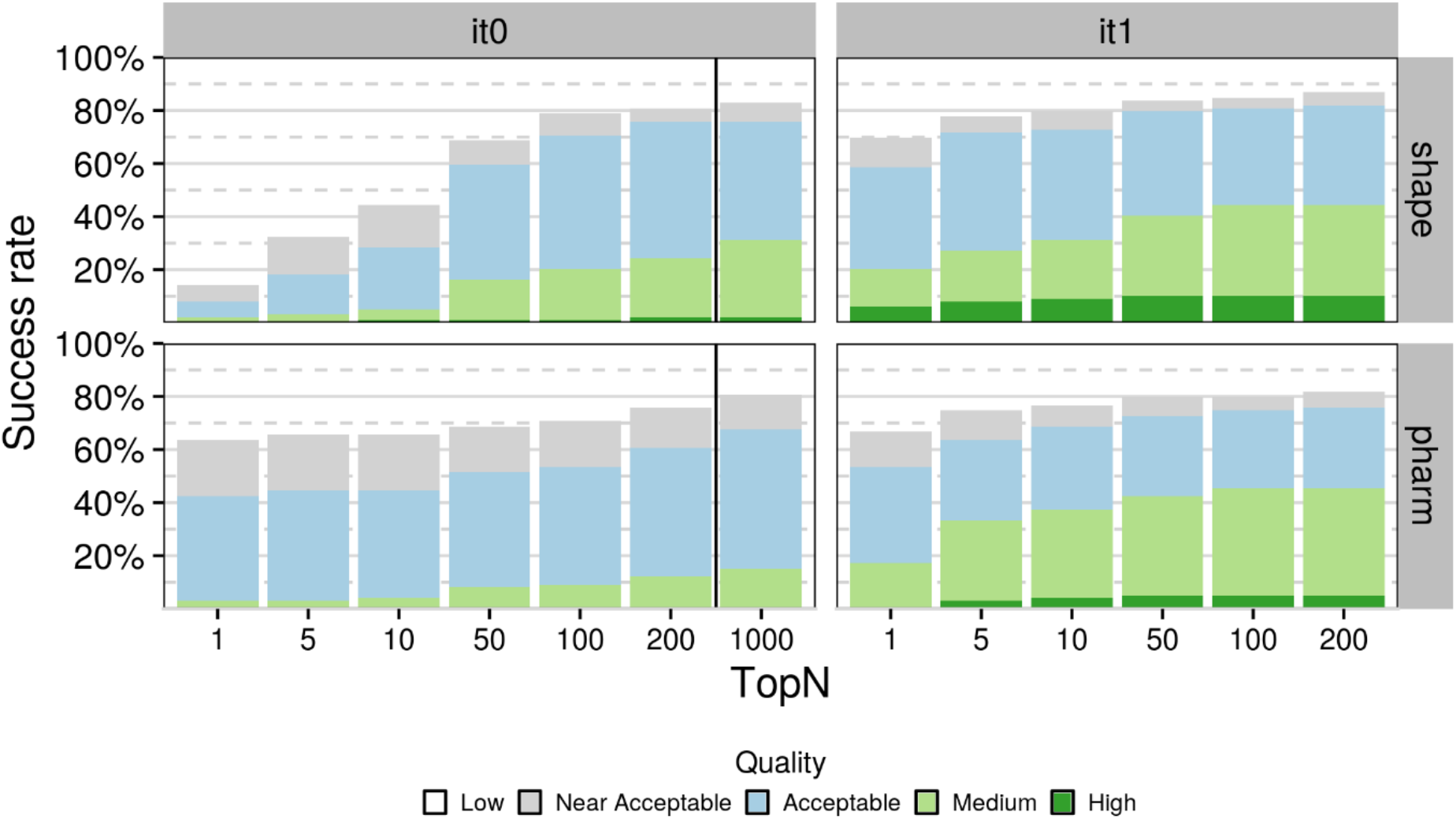
Success rate as a function of N-ranked models considered (TopN) for top 1, 5, 10, 50, 100, 200 and 1000 models for the initial rigid-body docking (it0) and after flexible refinement (it1). Each stacked bar plot corresponds to a specific cutoff with the colour of the bar indicating the distribution of model quality for that cutoff. Dark green, light green, light blue and light grey correspond to high-, medium-, acceptable- and near acceptable-quality models, respectively. The respective RMSD cutoffs are ≤ 0.5Å, ≤ 1Å, ≤ 2Å and ≤ 2.5Å when considering all heavy atoms of the compound after superimposing on the binding site backbone atoms of the receptors. Results have been grouped by protocol on the y axis and stage on the x axis. The black vertical line in the ‘it0’ indicates the cutoff at which models proceed to flexible refinement.

This pattern holds even if we include the “Near Acceptable” models – models whose RMSD to their respective receptor lies between 2 and 2.5Å – which cannot be considered near-native but are still likely to offer very relevant biological insights. When including these models, the respective percentages become 69.7, 77.8 and 86.7 for the shape top1, top5 and overall, and 66.7, 74.8 and 81.2 for the pharmacophore top1, top5 and overall. The performance of the pharmacophore protocol when considering only the top ranked model of it0 is worth mentioning as the success rate (when including the Near Acceptable models) stands at 63.6%, which makes it particularly impressive considering only rigid-body motions are allowed in it0. Despite this, the distribution of model quality also seems to favour the shape protocol as higher-quality models are generated for more targets at all stages and cutoffs.

The success rate, however, only tells one side of the story, as can be seen in Figure 3 which shows the distribution of model quality as a function of the rank for all targets for the refinement stage (Figure 3, bottom panel). Comparing the performance of the two protocols in this way reveals that the pharmacophore protocol produces models of acceptable (or better) quality much more consistently than the shape protocol but the shape protocol has higher coverage (see SI Figure 3 for rigid-body stage results). The top panel of Figure 3 shows the similarity between template and reference compounds (using the Tversky and Tanimoto metrics, respectively; see Methods for details) in bars and the 3D shape overlap in points. Comparing the performance of the two protocols with these metrics in mind reveals the 3D shape overlap to be a limiting factor for the outcome of the modelling, acting almost like a binary classifier, with targets whose overlap is below 0.5 almost never achieving good performance, whereas for those whose overlap is greater than 0.5 success is almost guaranteed, in particular for those whose overlap exceeds 0.75 (e.g., *adrb1* and *mcr*). Further comparing the performance of the two protocols, reveals that overlap is the feature than determines the outcome even when considering the same target. For example, for target *aces* the similarity and overlap values are 0.99 and 0.91, and 0.6 and 0.4, for the shape- and pharmacophore-based protocols, respectively, with the shape protocol yielding top-ranked, high-quality models and the pharmacophore-based protocol only producing Near Acceptable models despite a lower binding site RMSD for the template chosen for the pharmacophore-based protocol (0.09Å against 0.2Å for the shape-based protocol).

**Figure 3:**
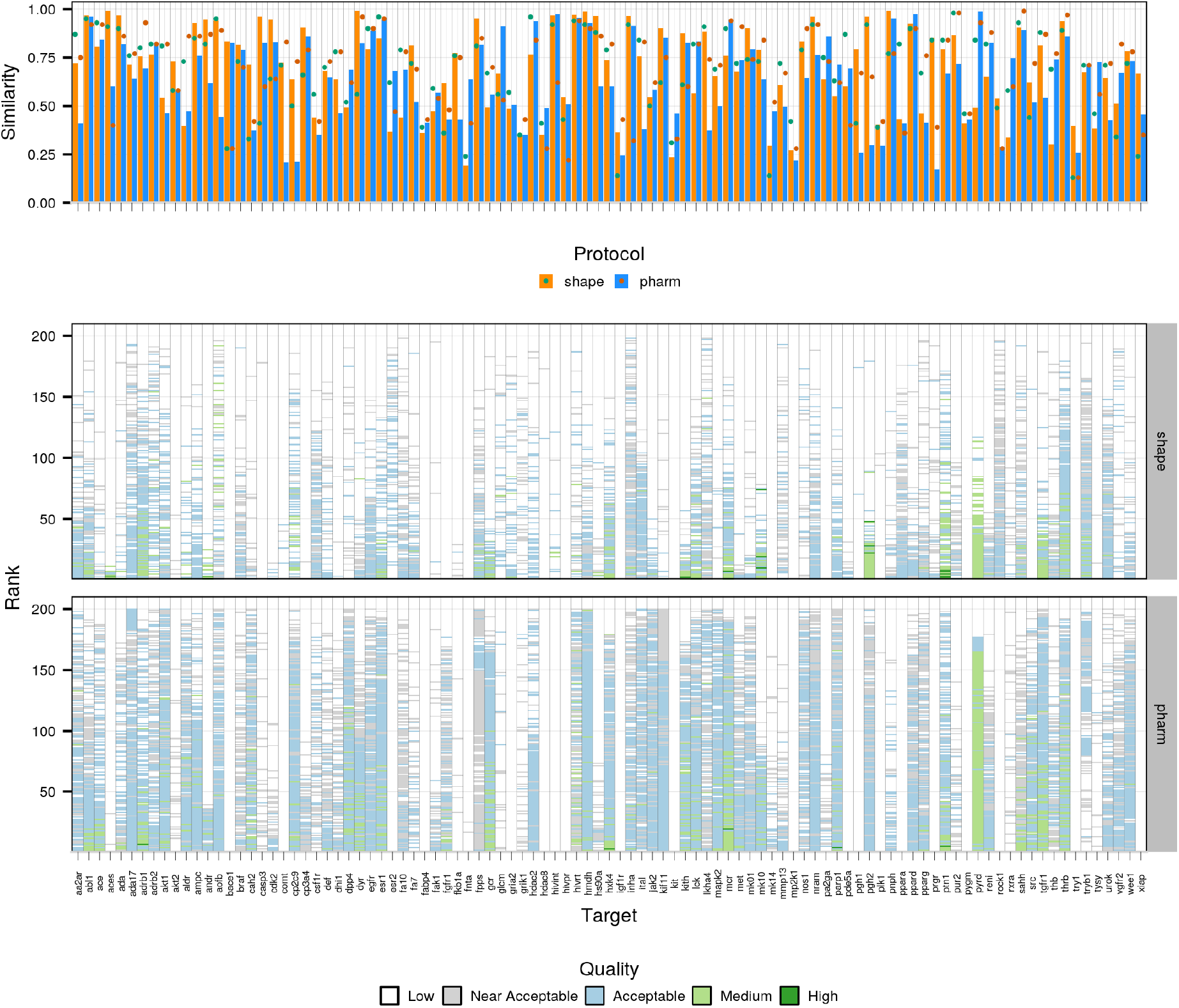
Comparison of template quality and performance for the two protocols. The top panel highlights the similarity (orange and blue bars for the shape- and pharmacophore-based protocols, respectively) and overlap (green and dark green dots for the shape- and pharmacophore-based protocols, respectively) between reference and template compounds. The bottom panel compares the performance for the semi-flexible refinement stage of HADDOCK (it1) for the two protocols. Each column corresponds to one target with the Y axis reflecting the ranking of models (ranks close to 0 refer to top-ranked models and those close to 200 to bottom-ranked models) and the colour of each model reflecting its quality (see description of Figure 2).

To further investigate the impact of template selection on the outcome of the modelling we compared the performance of the shape-based protocol in a subset of the dataset for which we identified both high-(similarity ≥ 0.8) and low-similarity templates (similarity around 0.4). Not only is the difference in terms of success rate stark, with the high-similarity subset observing success rates of 76.5%, 88.2% and 91.2% when considering top1, top5 and overall, respectively, against 44.1%, 47.1% and 73.5% for the low-similarity subset (still a very decent performance), but the simulations with the high-similarity templates clearly results in distributions that are more populated with higher-quality models (see SI Figure 4).

**Figure 4:**
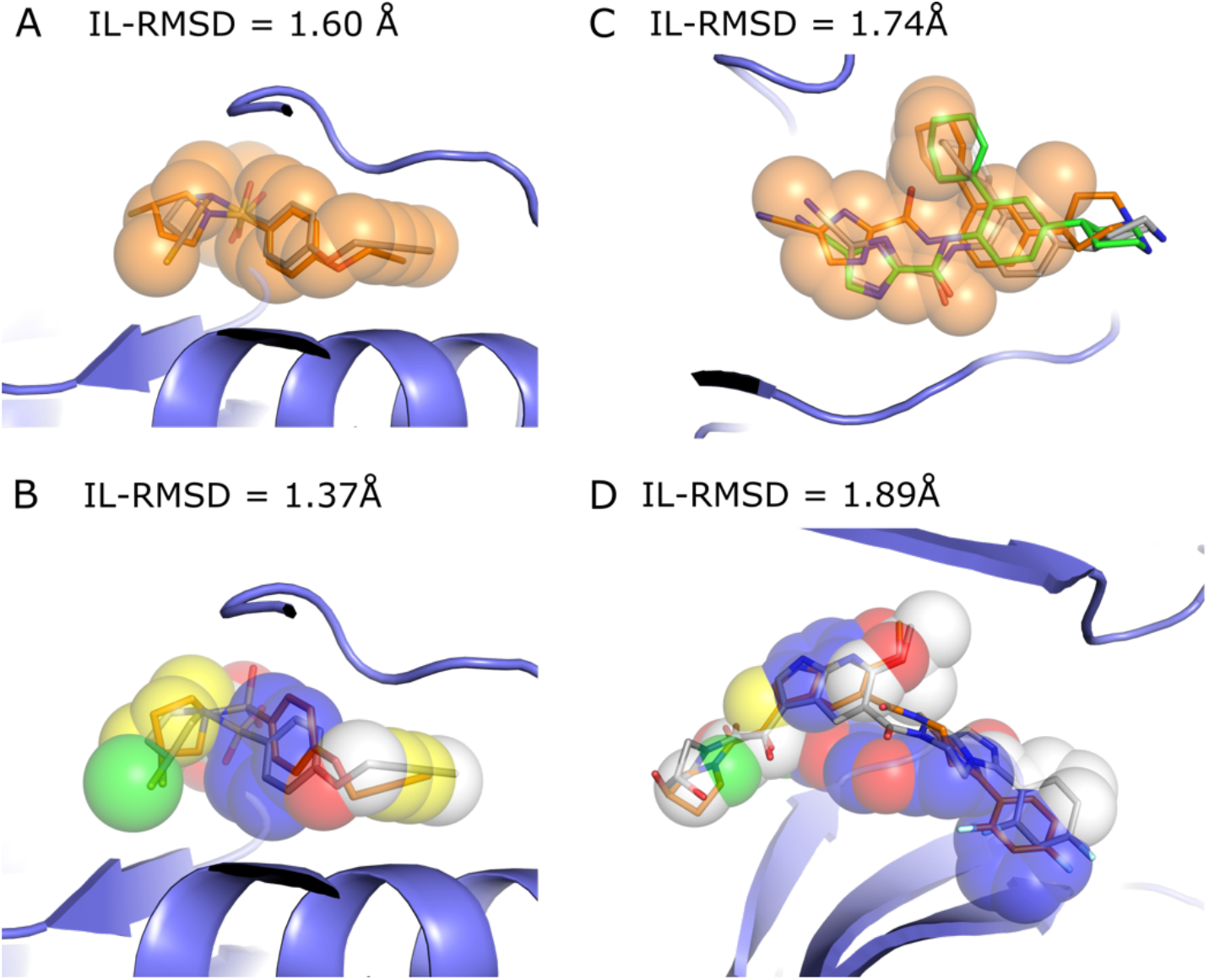
Illustration of shape-restrained docking outputs (orange sticks) given the input shape (spheres) as compared to the target binding mode (grey sticks). Panels **A** and **B** represent the top1 models generated by the shape- and the pharmacophore-based docking respectively for ada17 (ideal scenario), based on template 3b92. Panel **C** represents the best docking model generated in top5 (orange) and overall (green, with an IL-RMSD value of 1.74Å) for csf1r with the shape-based protocol, based on template 2i0v. Panel **D** represents the best docking model generated in top5 for mk14 with the pharmacophore-based protocol, based on template 3ha8. The colour code for the pharmacophore spheres is the following: H-bond donor (green), H-bond acceptor (red), hydrophobic (yellow), aromatic (blue) and regular shape bead (white)

Figure 4 A-B illustrates a docking scenario in which both shape-restrained protocols are expected to perform well due to the high similarity between the template and the target compounds (Tv = 0.714 and Tc = 0.641). Both protocols generate acceptable quality models at top1, with IL-RMSD (interface ligand-RMSD; see Methods for details) values of 1.60 Å and 1.37 Å for the shape- and the pharmacophore-based protocol respectively. Figure 4 C-D illustrates a more difficult scenario where the selected template differs substantially from the target compound (*csf1r* – Tv = 0.44 and *mk14* – Tc = 0.47, see Figure 6). We still observe good performance, with acceptable models generated at top2 for *csf1r* (1.74 Å) with the shape-based protocol, and at top5 for *mk14* (1.89 Å) with the pharmacophore-based protocol. This is mainly explained by a good shape overlap between the template and the target compounds (0.56 and 0.52).

#### Impact of the flexible refinement

As mentioned previously, we observe a significant improvement in the success rate at it1 as compared to it0 for the shape-based protocol, and a moderate improvement for the pharmacophore-based protocol. This underlines that the semi-flexible refinement plays a crucial role in improving the quality of the models generated at it0 as supported by Figure 5 (panel A). However, while we would expect moderate improvement related to local rearrangements in the binding site and relaxation of the small molecule, we also observe impressive and unexpectedly large improvement in the small molecule binding mode between it0 and it1, especially in the shape-based protocol. This can be clearly seen in the distribution of ΔIL-RMSDs for the shape protocol (orange distribution in Figure 5A) where the shoulder extends to up to 5Å (i.e. 5Å improvement) for the acceptable models after refinement. The distributions of the RMSD between the aligned template and reference small molecules, referred to as the ligand-ligand RMSD, show that both protocols also tend to generate models in which the conformation get closer to the reference for most targets (Figure S5). Three eloquent visual examples are provided in Figure 5B. The first one shows that the IL-RMSD of the top ranking it1 model for target *lck* (0.4 Å) is improved by 5.2 Å as compared to the corresponding model at it0. This target is considered an “easy case” since it is associated with an overlap of 0.82, and a binding site RMSD of 0.1 Å between the target and the template. The second example, an “intermediate case” (overlap of 0.57, binding site RMSD of 0.6 Å), shows that the IL-RMSD of the top5 it1 model for target *gria2* (1.0 Å) is improved by 4.1 Å. Finally, we present a “difficult case” with the target *pa2ga* for which no acceptable conformer was generated. Based on the excellent template that is selected by the pharmacophore-based protocol, the generated model ranked top10 at it1significantly improves on the ligand-ligand RMSD metric (Figure S5) as well as the IL-RMSD to the reference (it1 IL-RMSD = 1.8 Å with a ΔIL-RMSD = 3.9 Å). The ligand-ligand RMSD values are 0.35 Å, 0.76 Å and 1.72 Å for the three cases, respectively, indicating the excellent agreement in binding mode between our models and the reference structures.

**Figure 5.**
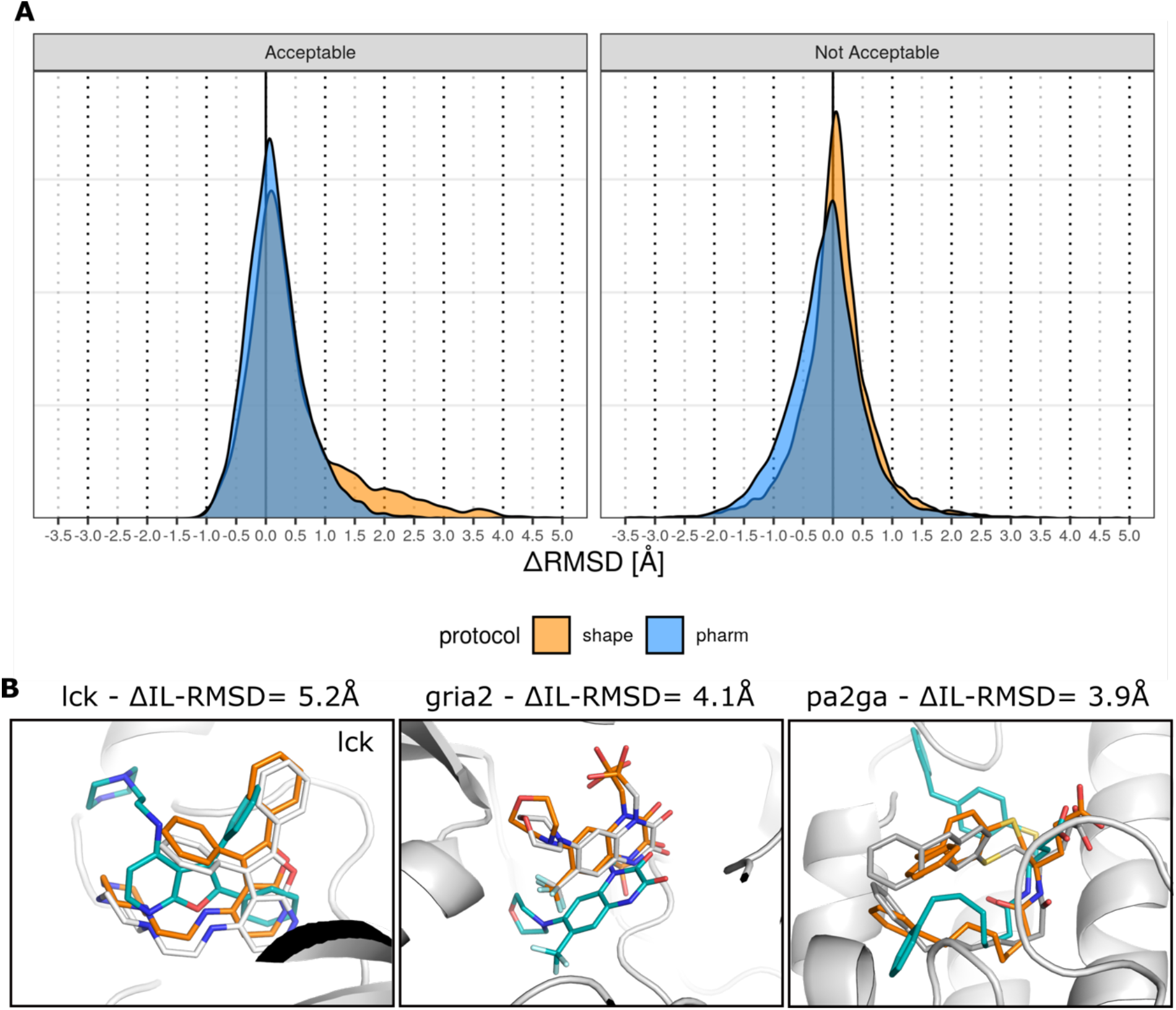
Illustration of the impact of the semi-flexible refinement on the docking models quality. Panel **A** shows the distribution of the Δ IL-RMSD between the models before and after semi-flexible refinement, computed as the IL-RMSD at it0 – the IL-RMSD at it1 for a given model. The figure is divided into two parts: the left sub-panel shows the distribution of Δ IL-RMSD for models that are acceptable at it1, the right sub-panel shows the distribution of Δ IL-RMSD for models that are not acceptable at it1. Positive values in the distribution indicate better it1 performance. Panel **B** illustrates three cases with large IL-RMSD improvement between the model generated at it0 (blue) and the refined model at it1 (orange) as compared to the bound conformation in the reference structure (grey). Templates 2of4, 3bki and 1kqu were used for cases lck, gria2 and pa2ga, respectively.

The overall larger improvement observed for the shape-based protocol as compared to the pharmacophore-based protocol could be explained by the higher degree of freedom provided by the dummy beads as compared to the pharmacophore-beads.

#### Peculiar cases

Since we always learn from errors, we investigated cases where shape-restrained docking fails. As the first HADDOCK step (it0) keeps both the receptor and the ligand rigid, it is essential to provide as starting point different conformers of the ligand to be docked. When the conformer generator fails in generating near-native conformations, the docking is likely to fail as well. Our results confirm this assumption as we observed that few to no acceptable models were generated in 7 cases with less than 10% acceptable conformers generated, (e.g. *cdk2*, *cp3a4*, *fkb1a*, *fnta*, *hivpr*, *pa2ga* and *xiap*). A second source or error is the binding site location of the template compound that may differ from the binding site of the target compound (Figure 6B.5), as highlighted by the Pearson correlation of 0.72 and 0.59 observed between the distance in geometric centre of the template and the target compounds and the best IL-RMSD obtained with the shape- and pharmacophore-based protocol respectively (Figure S6). External expert-eye or experimental knowledge should alleviate this issue and help filtering out potential erroneous template compounds in early stages of the protocol. For this work we however followed an automated pipeline. Third, if both protocols are capable of generating good quality models where the template compounds show poor similarity with the target compounds (Figure 6A), the opposite also happens. In some cases, our initial assumption stating that similar compounds that bind the same target receptor should in principle adopt similar binding modes is not validated. This is the case for the *hivint* and *bace1* target and template compounds as illustrated on Figure 6B. The different structures of *hivint* complexed with N-benzyl indolin-2-ones reported in the original paper of the target structure [27] display various binding modes of the compounds in the same binding site. In the case of *bace1*, two binding modes of iminohydantoin bace1 inhibitors are reported with a 180° flip of the iminohydantoin core [28]. Finally, a visual inspection could beneficially help in discarding template compounds that share poor similarity with the target compound in terms of shape, flexibility and physico-chemical properties despite a meaningful similarity score (e.g., Figure 6B.5).

**Figure 6:**
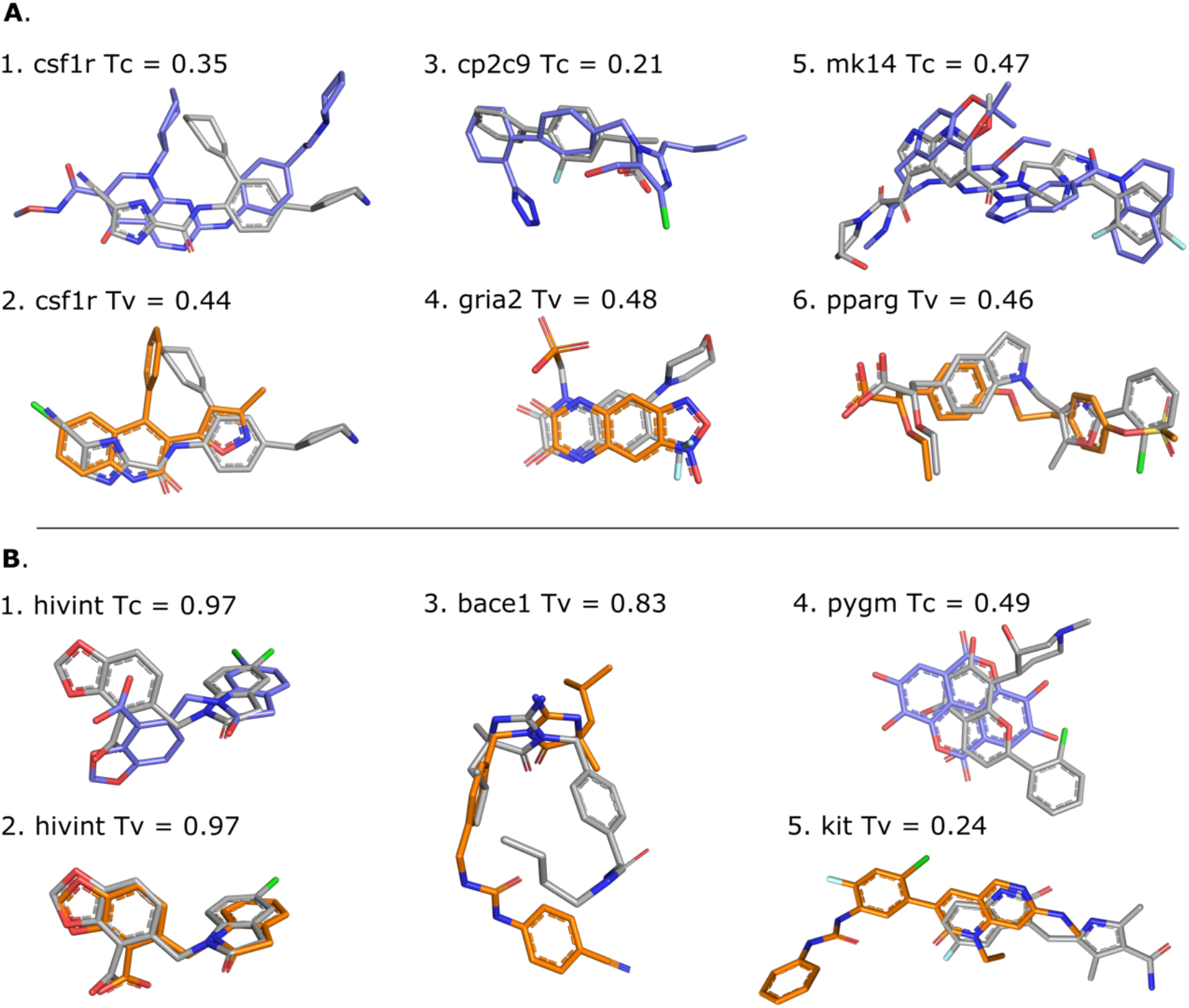
Illustration of template (shape-based protocol: orange, pharmacophore-based protocol: blue) and target (grey) compounds. **Panel A** shows different template/target compound couples of low similarity, i.e. Tanimoto (Tc) or Tversky (Tv) coefficient < 0.5, that lead to successful docking. Templates 3dpk, 2i0v, 5×23, 3bki, 3ha8 and 1i7i were used for cases 1-6, respectively. **Panel B** show scenarios that lead to docking failure for different possible reasons. While displaying high similarity with the target compounds, the hivint pharmacophore-based template compound adopts a different binding mode (**B.1**), and the shape-based template compounds leads to an unexpected failure that cannot be explained neither by the coverage nor their respective binding site (**B.2**). The shape-based template compound for bace1, that is the same as the pharmacophore-based template, adopts a different binding mode than the target despite the high similarity (**B.3**). Low shape, flexibility and physico-chemical similarities are observed between the template and the target compound for pygm (**B.4**); The case **B.5** is an example of poor overlap between template and target compounds that is responsible for poor performance. Templates 3nf9, 3nf6, 3l5c, 3g72 and 6mob were used for cases 1-5, respectively.

#### Which protocol should be favoured?

The shape-based protocol yields better performances in terms of coverage, i.e. it generates near-native models, including near-acceptable models, for more DUD-E complexes (top1: 69.7%, top5: 77.8 %, overall: 87%) than the pharmacophore-based protocol (top1: 66.7%, top5: 74.8 %, overall: 81.2 %). In an ideal scenario where we merge the results of the two protocols and pick up the best model (i.e. lowest IL-RMSD) per rank, the global coverage is remarkably improved (top1: 81%, top5: 86%, overall: 91%). As illustrated in Fig 2, it means that even though the protocols display comparable performance on most targets, one protocol could be favoured over the other in some cases (e.g. *aces*, *fkb1a*, *gria2*, *hivint*, *mapk1*, *mk14*, *pde5a*, *prgr*). We identified the overlap between the template and the target compounds shapes as the determinant indicator of the success of the shape- and the pharmacophore-based docking (Figure S7). The overlap does not only inform on the proximity between the binding sites of the template and the target compounds, it also tells about the binding mode similarity. However, this information is accessible only in benchmarking conditions where the crystallographic binding mode of the target ligand is known. To mimic real-life conditions, we investigated if different combinations of descriptor filters extracted from the unbound form of the target and the template compounds (similarity, difference in molecular weight, number of rotatable bounds and accessible surface area) could lead to an enrichment in the success-rate. We identified no evident conditions to favour one protocol over the other, but it is worth mentioning that both protocols achieve high success rates when the template and target share moderate to high similarity (≥~0.4/0.3 for shape and pharmacophore, respectively) (Figure 7). As such, the shape-based protocol performance (including near-acceptable models) rises to 74.1 %, 83.5 % and 92.9 % at top1, top5 and overall (85 targets), when considering a Tversky similarity ≥ 0.4. With similar conditions, i.e. a Tanimoto coefficient over 0.3, the performance stands at 71.9 %, 80.9 % and 87.6 % at top1, top5 and overall (89 targets) with the pharmacophore-based protocol. Adding a molecular weight filter (|MW_ref_ – MW_temp_| < 50 g/mol) slightly improves the performance which rises to 77.5 %, 89.8 % for the shape top1 and top5, and reaches an impressive overall performance of 100% when considering all generated models. A lower overall performance is reached with the pharmacophore-based protocol (93.2 %), but we observe a higher early enrichment of 81.4 % at top1. Of note, 49 and 59 DUD-E targets fulfil these two criteria for the shape- and the pharmacophore-based protocol, respectively. All the aforementioned analyses suggest the shape-based protocol should be favoured for the sake of coverage. However, the conditions we identified could be used to favour one of these protocols depending on the available data and the intended goal of the study.

**Figure 7:**
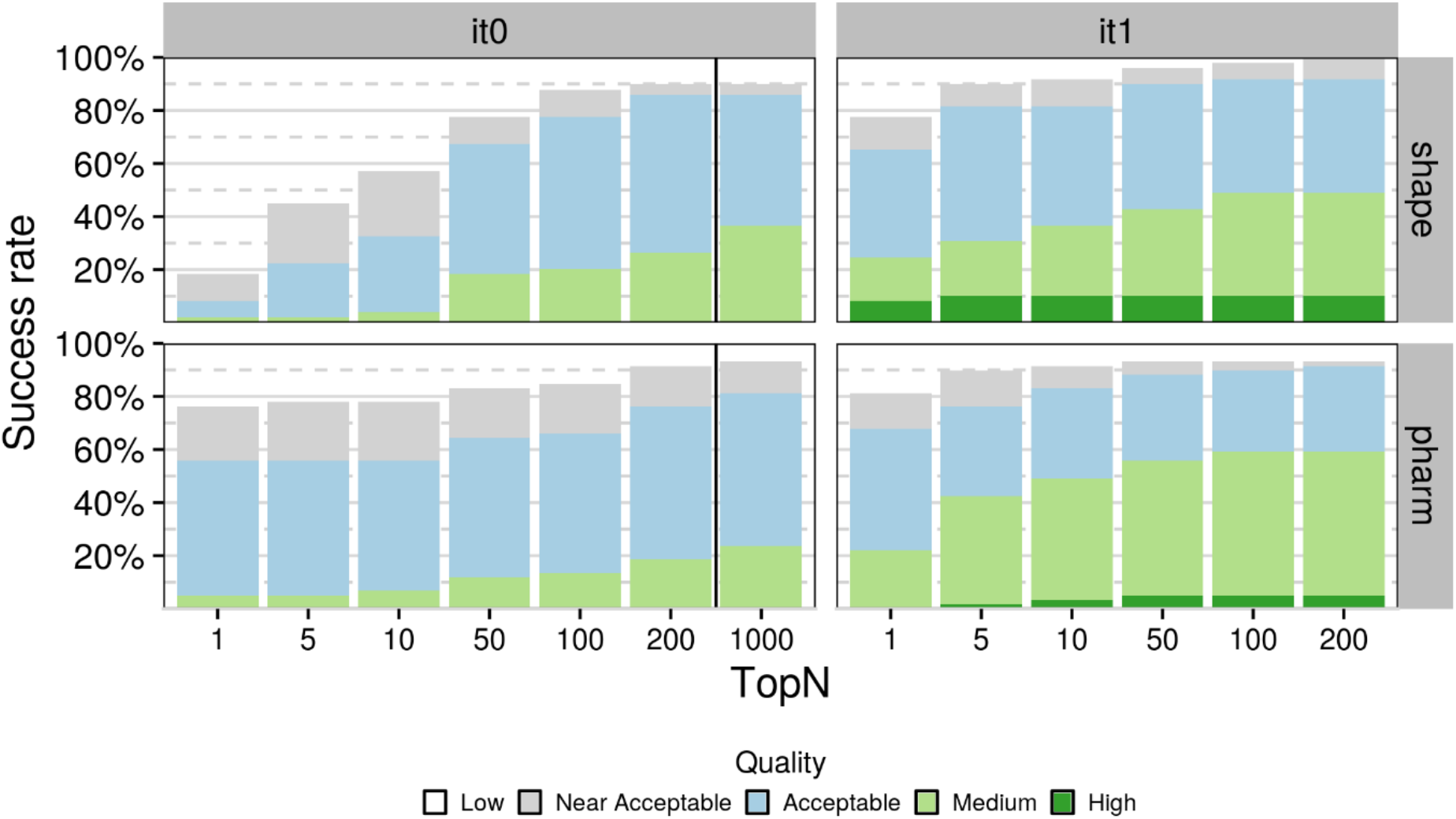
Success rate of the shape- and pharmacophore-based protocol on DUD-E complexes (49 and 59, respectively) that match two criteria: 1) the similarity coefficient between template and the reference compounds is higher than 0.4 and 0.3, respectively, and 2) the difference in heavy atom molecular weight is below 50g.mol^−1^.

#### Comparison to state-of-the-art docking tools

A similar benchmarking study was conducted with four commercial docking programs, namely Gold [29], Glide [30–32], Surflex [33]and FlexX [34], which are all widely used in industry and have better sampling power than most non-commercial docking programs [35]. In this study, two of the 102 DUD-E targets (*aofb* and *casp3*) were discarded as they show covalent binding to their respective receptor. Of note, those two targets are part of our dataset and the three we are missing are part of their dataset. To fairly compare the performance, we discarded *aofb* and *casp3* from our results. Commercial programs were evaluated in bound-unbound docking mode, i.e. unbound conformations of the compounds have been docked onto the bound conformation of the receptor (i.e. no conformational changes required on the protein side), while we did perform fully unbound docking which represents a significantly more difficult – yet more realistic – scenario [11]. Our two protocols show very competitive overall performance with those commercial docking programs, the shape-based protocol ranking 2^nd^ with a success rate of 81 % just behind Surflex (84 %) (Figure S8), and the pharmacophore-based protocol ranking 5^th^ with 74 %. We also observe rather competitive early enrichment with 54 % and 58 % success at top1 for the fully unbound pharmacophore- and the shape-based protocols respectively against 64 % and 65 % for Gold and Glide respectively. It is interesting to note that all commercial software fail to generate near-acceptable models for 6 DUD-E targets on which at least one of our protocols also fails (*bace1*, *cdk2*, *fkb1a*, *fnta*, *hivint*, *pgh1*) for different reasons (see *Peculiar Cases*).

## Conclusion and Perspectives

We have presented two new protocols for the modelling of protein-small molecule complexes with HADDOCK, which make excellent use of the integrative modelling capabilities of the platform, to yield very promising results. Selection of suitable templates for the modelling based on ligand similarity and use of shape data during the docking lie at the core of both approaches. In the first, the shape is only defined in geometrical terms whereas in the second it is additionally annotated with pharmacophore data, thus naming the two protocols shape- and pharmacophore-based. The dataset on which these protocols were benchmarked is based on the widely used DUD-E dataset, which was extended to make use of unbound receptors (or rather receptors bound to another ligand, which is commonly defined as cross-docking). This “unbound” DUD-E dataset, which is freely available in ready-to-dock format on GitHub, is composed of diverse targets of various difficulty degrees. Our analysis has revealed that the main limiting factor for the outcome of the docking with our shape protocols is the 3D overlap between template and reference compounds, in other words, whether the template and reference compounds not only occupy the same binding pocket but have similar binding modes. We have also shown that we can predict the reliability of the modelling based on features that can be computed in real-world usage: The difference in molecular weight between template and reference compounds and their molecular similarity. When we select the subset of our dataset for which the compounds are not too dissimilar in terms of molecular similarity (Tv and Tc greater or equal than 0.4 and 0.3 for the shape- and pharmacophore-based protocols, respectively) and molecular weight (|MW_ref_ – MW_temp_| < 50 g/mol) we achieve success rates of 65.3%, 81.6% and 91.8%, and 67.8%, 76.2% and 91.5% when considering top1, top5 and overall, for the shape- and pharmacophore-based protocols, respectively. Using all targets, the performance remains rather high with 58.5%, 71.7% and 81.8%, and 53.5%, 63.6% and 75.8% for the two protocols. The performance of our protocols is on par with that of leading small molecule docking software, despite the fact our approach was benchmarked on a fully unbound dataset. As such it is representative of the performance one can expect when using this pipeline in a real-life scenario where one would not have access to the bound form of the receptor and/or that of the compound.

Despite this promising performance our method still suffers from a few limiting factors. Chief among those is the dependency on a template whose compound occupies the same binding pose as that of the ligand to be modelled for a successful prediction. This was identified as the most severe limiting factor for both protocols. It is a problem which is not unique to these protocols but is shared by all data-driven methods, HADDOCK being no exception [36]. Overcoming this obstacle is no easy task and will require the development of methodology for the automated identification, combination and weighting of multiple template shapes and receptors in order to increase the coverage of the binding site. Another area where these protocols could benefit from further improvements is the ranking of solutions during the flexible stage as can be seen by the difference in success rate at the top1 or top5 level and overall. Also worth noting is the fact that, although regular users of the command-line version of HADDOCK will be able to make use of this pipeline, it might be well out of reach for most researchers who have no expertise in programming and the use of computational chemistry toolkits. Therefore, it would be of interest if these protocols were to be made available through the freely accessible HADDOCK webserver (https://wenmr.science.uu.nl/haddock2.4/). We hope to expand on these three points in the near future.

Limitations and future improvements mentioned in the previous paragraph notwithstanding, the protocols we are proposing achieve high performance on experimental conditions that mimic real-life ones as closely as possible using unbound structures for receptors as well as compounds. Furthermore, the performance we are reporting is on par with that of leading commercial docking platforms when benchmarked on partially bound datasets, and in many cases exceeding it, thus making it an excellent option for the modelling of protein-small molecule complexes.

## Materials and Methods

### Dataset

To validate our new approach, we decided to benchmark the performance of these protocols against the Enhanced Database of Useful Decoys dataset (DUD-E – http://dude.docking.org/), one of the most widely used small-molecule docking benchmarks [20]. In total, the DUD-E consists of 102 targets with each target corresponding to a single protein receptor (usually of pharmaceutical interest) with a compound bound to it. The Protein Data Bank (PDB) [37] entries associated with every DUD-E target became our reference receptors. We discarded three targets (*cxcr4*, *drd3*, *kpcb*) because we could not identify any viable templates for them (see next section) bringing our total to 99 targets. The structures of the reference receptors were downloaded from the PDB instead of the DUD-E website to avoid any post-processing that might have been applied to them.

### Template Identification

The procedure we followed for identifying a template for each of the 99 targets in our dataset can be summarised in the following steps:

1. Search the PDB for appropriate template structures. We used two approaches in conjunction with a few criteria to identify templates:
  a. Search for receptors whose sequence was more than a predetermined cutoff similar to the sequence of our reference receptor. The precise value we used for the sequence similarity cutoff differed from target to target because of the intrinsic features of some targets. As an example, some targets were membrane receptors that had been co-crystallised with a lysozyme domain. The presence of the lysozyme sequence, lysozyme being one of the most commonly encountered domains in the PDB, affected the results and we had to take that into account when adjusting the sequence similarity cutoff. The default value was 90% and in cases such as the one mentioned above was adjusted after manual inspection of the results.
  b. Search for receptors bound to compounds which were similar to the reference compound. For this we made use of the “Fingerprint Similarity” feature of the “Chemical” tab of the Advanced search functionality of the RCSB portal of the PDB (https://www.rcsb.org/search/advanced).
2. After pooling the results from both searches, we iteratively removed unsuitable templates based on a few criteria:
  a. Templates which contained only irrelevant compounds such as crystallisation buffer artefacts or some of the most commonly encountered compounds such as hemes.
  b. Incorrect templates based on sequence identity (regarding the templates identified through ligand similarity).
  c. Templates whose compounds were found to be stereoisomers of the reference compounds.
3. After removal of all unsuitable templates, we calculated the pairwise similarity between all reference and template compounds.
  a. *Shape-based protocol*: The similarity metric we chose for this is the Tversky similarity [38] (with weights for the reference and template compounds set to 0.8 and 0.2, respectively) [39,40] computed over the Maximum Common Substructure (MCS) as identified with the rdFMCS implementation of RDKit (version 2020.09.3) [41].
  b. *Pharmacophore-based protocol*: We computed the Tanimoto coefficient over 2D pharmacophore fingerprints generated with RDKIT [41] using the default pharmacophore fingerprint factory.
4. After calculating all similarities, we rank the templates according to the similarity of their compound to the reference compound and select the one with the highest similarity with the condition that the template and the target share the same binding site (defined as < 5 Å around the bound small molecule), i.e. 100% sequence identify in the binding site.

### Conformer generation

To ensure there is no bias during the docking, instead of starting from compounds bound to the reference – or other – receptors, we generate 3D conformations of the target compounds starting from their isomeric SMILES [42,43] with RDKit using the 2020 parameters (only the small aliphatic ring subset) with energy minimisation and the ETKDG algorithm [44,45]. We cap the maximum number of conformers to 50 and provide the ensemble of conformers to HADDOCK for docking.

### Docking

#### System Preparation

Prior to docking we process the structures to ensure the residue numbering between template and reference structures is consistent (a necessity for the analysis), rename residues, cofactors and ions according to HADDOCK specifications, remove nonbiological symmetric units and remove all irrelevant artefacts such as crystallisation buffers. We generated topology files of the compounds to be docked using PRODRG (version 070118.0614).

#### Shape-based protocol

After identifying one receptor template per target, we transform all heavy atoms of its compound into dummy beads (those do not interact with the remaining of the system). We then define ambiguous distance restraints with an upper limit of 1Å between the shape beads and the heavy atoms of the compound to be docked. The nature of the restraints creates an additional consideration, specifically what should be the “origin” and “target” of the restraints? In this protocol, the directionality of the restraints depends on the size of the reference and template compounds, in terms of heavy atom count. Specifically, the restraints are always defined from smaller to larger. If the reference compound is larger than the template compound, the restraints are defined from each shape bead to any compound heavy atom. If the opposite is true (the template compound is larger than the reference compound) then restraints are defined between each heavy atom of the conformer to any shape bead. These ambiguous distance restraints effectively enforce that ligand atoms and beads must overlap. Depending on the directionality of the restraints part of the ligand might remain outside the shape defined by the beads and vice-versa. All shape restraints are used during the simulation.

#### Pharmacophore-based protocol

In addition to the transformation of the template atoms into dummy beads, we assigned pharmacophore features to the beads to add physico-chemical properties to the spatial information they hold. We assigned a pharmacophore features (H-bond donor, H-bond acceptor, negative ionizable, positive ionizable, zinc binder, aromatic, hydrophobic and lumped hydrophobic) to each atom with RDKIT using the default SMARTS-based feature definition. If no pharmacophore feature is assigned to an atom, a regular shape bead is defined. We defined distance restraints with an upper limit of 1Å between the beads and the atoms of the compounds to be docked provided that they share the same pharmacophore feature – or no pharmacophore feature. In this protocol, we cannot impose restraints from the smaller to the larger anymore because restraints are defined between features of the same class -- some classes may be more populated in the reference compound while others may be more populated in the template compound. Instead, we defined all restraints from the compound to the beads. Similarly to the shape-based protocol, all restraints are considered during the docking process.

Fig. 1 illustrates the shape-based and the pharmacophore-based docking protocol.

#### HADDOCK

For the docking we use the January 2021 release of the command-line version of HADDOCK 2.4. The number of models generated in the initial rigid-body docking stage (it0) of HADDOCK is set to 20 times the number of starting ligand conformations. This sampling ratio was found to be the most performant during benchmarking (data not shown). If the number of it0 models that are sampled is larger than 200 then only the top 200 it0 models proceeded to flexible refinement, otherwise all models do. We only perform the semi-flexible refinement stage of HADDOCK (it1) and skip full water refinement (itw) as this final stage is not improving the results as already remarked from our D3R participation. The positions of both the receptor and its associated shape are fixed in their original orientations while the ligand is translated away from the protein and randomly rotated for each docking trial. The shape is kept rigid throughout the protocol while the receptor interface and the ligand become flexible during the refinement stage. Systematic sampling of 180° rotations along the interface is disabled for it0. We also scale down the intermolecular interactions during the rigid body stage to facilitate the insertion of the ligand into the binding pocket and accordingly exclude the vdW energy term during the scoring of the rigid-body models. The modified parameter settings for HADDOCK are summarized in Table S3.

Other than the above defined modifications, the scoring function used is the default scoring function of HADDOCK. Its functional form, specific for protein-ligand docking for the two stages is:

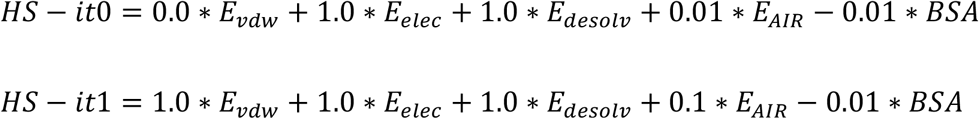

where E_vdw_, E_elec_ and E_desolv_ stand for van der Waals, Coulomb electrostatics and desolvation energies, respectively and BSA for the buried surface area. The non-bonded components of the score (E_vdw_, E_elec_) are calculated with the OPLS forcefield [46]. The desolvation energy is a solvent accessible surface area-dependent empirical term [47] which estimates the energetic gain or penalty of burying specific sidechains upon complex formation.

### Evaluation of results

We evaluate the quality of the generated models according to their structural deviation from the reference structures. For this we use the interface-ligand RMSD (IL-RMSD), which is the RMSD calculated over all heavy atoms of the ligand after superimposing on all backbone atoms of the interface of the receptor. Models with an IL-RMSD of less than 0.5 Å, between 0.5 and 1 Å, between 1 and 2 Å, between 2 and 2.5 Å and over 2.5 Å are classified as high-, medium-, acceptable-, near acceptable- and low-quality, respectively. The initial fitting was performed using the McLachlan algorithm [48] as implemented in the program ProFit (Martin, A.C.R., http://www.bioinf.org.uk/software/profit/ - available through SBGrid [49]). Calculation of symmetry-corrected RMSD values for the compounds were performed with *obrms* from the Open Babel distribution (https://github.com/openbabel/openbabel-version3.1.1) [50]. As part of the analysis and interpretation of the results we also present a classification of target difficulty based on the 3D shape overlap of target and reference compounds after fitting on the binding site backbone atoms of their respective receptors. Fitting was performed with ProFit using the same settings as previously mentioned and the 3D shape overlap was calculated with the *Exact Overlap* metric of the shape toolkit of OpenEye (release 2020.2.0) [51–53].

### Data availability

All docking models generated during the benchmarking of the two protocols with HADDOCK 2.4 are made available through our laboratory data collection at https://data.sbgrid.org/labs/32/ [54] (*Note for reviewers: The full dataset will be deposited upon acceptance*). The unbound DUD-E benchmark data set, with all the code, docking input and parameter files, results and analysis files are made available through GitHub (https://github.com/haddocking/shape-restrained-haddocking).

## Acknowledgments

We would like to acknowledge support from the European Union Horizon 2020 projects BioExcel (675728, 823830) and EOSC-hub (777536) and from the Innovative Medicines Initiative 2 Joint Undertaking (JU) under grant agreement No 101005077. The JU receives support from the European Union’s Horizon 2020 research and innovation programme, EFPIA, BILL & MELINDA GATES FOUNDATION, GLOBAL HEALTH DRUG DISCOVERY INSTITUTE and UNIVERSITY OF DUNDEE.

## Abbreviations Used

HADDOCK: High Ambiguity Driven DOCKing
DUD-E: Database of Useful Decoys: Enhanced
GC: Grand Challenge
D3R: Drug Design Data Resource
CSAR: Community Structure-Activity Resource
IL-RMSD: Interface Ligand-RMSD
ΔIL-RMSD: delta IL-RMSD
MCS: Maximum Common Substructure

