## Supplementary material for "Shape-restrained modelling of protein-small molecule complexes with HADDOCK"

### Table of Contents

**Table S1** .................... **S3**

**Table S2** .................... **S6**

**Table S3** .................... **S9**

**Figure S1** .................... **S10**

**Figure S2** .................... **S11**

**Figure S3** .................... **S12**

**Figure S4** .................... **S13**

**Figure S5** .................... **S14**

**Figure S6** .................... **S15**

**Figure S7** .................... **S16**

**Figure S8** .................... **S17**

***Table S1**: Templates used for the shape-based protocol. The first column lists the DUD-E target according to DUD-E nomenclature, the second column lists the PDB id of the identified template, the third column lists the RMSD between template and reference receptors using the backbone atoms of the binding site (all residues whose atoms lie within 5Å of the reference compound), the fourth through sixth columns list the interface ligand RMSD (heavy-atom RMSD between reference and model compound after superimposing on backbone atoms of the receptor binding site) when considering top1, top5 and all models generated during the semi-flexible refinement stage.*

| TARGET | TARGET PDBID | TEMPLATE PDBID | BINDING SITE RMSD [Å] | TOP1 [Å] | TOP5 [Å] | BEST [Å] |
|---|---|---|---|---|---|---|
| **AA2AR** | 3eml | 5iu8 | 0.24 | 1.46 | 1.18 | 0.90 |
| **ABL1** | 2hzi | 1opk | 0.26 | 0.91 | 0.51 | 0.51 |
| **ACE** | 3bkl | 3bkk | 0.09 | 1.58 | 1.58 | 1.40 |
| **ACES** | 1e66 | 6g1v | 0.21 | 0.44 | 0.43 | 0.43 |
| **ADA** | 2e1w | 1ndy | 0.29 | 0.46 | 0.46 | 0.46 |
| **ADA17** | 2oi0 | 3b92 | 0.57 | 1.60 | 1.60 | 1.22 |
| **ADRB1** | 2vt4 | 2ycz | 0.25 | 0.83 | 0.74 | 0.59 |
| **ADRB2** | 3ny8 | 3nya | 0.37 | 2.54 | 0.96 | 0.74 |
| **AKT1** | 3cqw | 3mv5 | 0.40 | 1.84 | 1.06 | 0.91 |
| **AKT2** | 3d0e | 3e88 | 0.43 | 2.95 | 2.47 | 2.26 |
| **ALDR** | 2hv5 | 2pdg | 0.42 | 0.67 | 0.59 | 0.59 |
| **AMPC** | 1l2s | 1xgi | 0.21 | 1.25 | 1.19 | 0.56 |
| **ANDR** | 2am9 | 1gs4 | 0.24 | 0.44 | 0.44 | 0.44 |
| **AOFB** | 1s3b | 2c75 | 0.08 | 1.60 | 1.26 | 0.62 |
| **BACE1** | 3l5d | 3l5c | 0.11 | 5.45 | 5.28 | 3.32 |
| **BRAF** | 3d4q | 3psd | 0.34 | 1.36 | 1.01 | 1.01 |
| **CAH2** | 1bcd | 6rzx | 0.12 | 1.23 | 1.23 | 1.23 |
| **CASP3** | 2cnk | 2c2m | 0.13 | 4.10 | 3.01 | 2.18 |
| **CDK2** | 1h00 | 1v1k | 0.46 | 1.69 | 1.69 | 1.69 |
| **COMT** | 3bwm | 3s68 | 0.15 | 4.31 | 3.48 | 1.07 |
| **CP2C9** | 1r9o | 5x23 | 0.74 | 6.64 | 1.88 | 0.70 |
| **CP3A4** | 3nxu | 4i4g | 0.75 | 2.15 | 2.15 | 2.15 |
| **CSF1R** | 3krj | 2i0v | 0.34 | 2.17 | 1.74 | 1.03 |
| **DEF** | 1lru | 3k6l | 0.51 | 1.33 | 1.33 | 1.32 |
| **DHI1** | 3frj | 3pdj | 0.86 | 7.99 | 1.39 | 1.39 |
| **DPP4** | 2i78 | 1x70 | 0.23 | 2.86 | 2.57 | 1.70 |
| **DYR** | 3nxo | 3nxy | 0.44 | 1.51 | 1.34 | 1.00 |
| **EGFR** | 2rgp | 3bel | 0.11 | 1.40 | 1.04 | 1.04 |
| **ESR1** | 1sj0 | 1xpc | 0.23 | 1.04 | 0.77 | 0.72 |
| **ESR2** | 2fsz | 1qkn | 0.46 | 1.44 | 1.44 | 1.02 |
| **FA10** | 3kl6 | 2cji | 0.34 | 1.35 | 1.24 | 1.24 |
| **FA7** | 1w7x | 1w2k | 0.51 | 1.73 | 1.48 | 0.94 |
| **FABP4** | 2nnq | 6ljs | 0.37 | 7.41 | 3.84 | 2.92 |
| **FAK1** | 3bz3 | 6i8z | 0.29 | 3.04 | 2.49 | 1.25 |

| | | | | | | |
|---|---|---|---|---|---|---|
| **FGFR1** | 3c4f | 1agw | 2.12 | 7.19 | 3.55 | 3.55 |
| **FKB1A** | 1j4h | 1j4i | 0.44 | 6.15 | 2.08 | 1.85 |
| **FNTA** | 3e37 | 1x81 | 0.51 | 9.84 | 9.36 | 4.43 |
| **FPPS** | 1zw5 | 4ga3 | 0.20 | 0.91 | 0.86 | 0.63 |
| **GCR** | 3bqd | 1m2z | 0.54 | 0.71 | 0.58 | 0.58 |
| **GLCM** | 2v3f | 6tjk | 0.31 | 3.98 | 3.08 | 0.98 |
| **GRIA2** | 3kgc | 3bki | 0.60 | 1.28 | 0.98 | 0.98 |
| **GRIK1** | 1vso | 2wky | 1.97 | 2.06 | 1.76 | 1.70 |
| **HDAC2** | 3max | 5iwg | 0.12 | 7.31 | 1.22 | 1.08 |
| **HDAC8** | 3f07 | 5bwz | 0.66 | 8.26 | 3.73 | 1.89 |
| **HIVINT** | 3nf7 | 3nf6 | 0.21 | 1.92 | 1.92 | 0.66 |
| **HIVPR** | 1xl2 | 3bhe | 0.22 | 4.73 | 3.51 | 2.82 |
| **HIVRT** | 3lan | 3lam | 0.38 | 1.65 | 1.26 | 0.81 |
| **HMDH** | 3ccw | 3ccz | 0.13 | 2.05 | 1.41 | 1.03 |
| **HS90A** | 1uyg | 1uyh | 0.14 | 0.80 | 0.80 | 0.80 |
| **HXK4** | 3f9m | 3fr0 | 0.94 | 0.83 | 0.69 | 0.69 |
| **IGF1R** | 2oj9 | 3o23 | 2.94 | 11.60 | 10.84 | 4.94 |
| **INHA** | 2h7l | 4u0j | 0.17 | 2.07 | 0.69 | 0.69 |
| **ITAL** | 2ica | 3m6f | 0.27 | 1.80 | 1.45 | 0.94 |
| **JAK2** | 3lpb | 3e64 | 0.81 | 1.67 | 0.93 | 0.93 |
| **KIF11** | 3cjo | 2fky | 0.31 | 1.02 | 1.02 | 1.02 |
| **KIT** | 3g0e | 6mob | 0.46 | 8.64 | 6.87 | 3.16 |
| **KITH** | 2b8t | 2uz3 | 0.20 | 0.60 | 0.45 | 0.45 |
| **LCK** | 2of2 | 2of4 | 0.11 | 0.41 | 0.41 | 0.41 |
| **LKHA4** | 3chp | 3chr | 0.15 | 2.31 | 2.00 | 0.82 |
| **MAPK2** | 3m2w | 3m42 | 0.66 | 7.42 | 7.25 | 1.14 |
| **MCR** | 2aa2 | 4uda | 0.26 | 0.45 | 0.45 | 0.45 |
| **MET** | 3lq8 | 5dg5 | 0.93 | 1.00 | 1.00 | 1.00 |
| **MK01** | 2ojg | 2oji | 0.95 | 1.72 | 1.24 | 0.90 |
| **MK10** | 2zdt | 2zdu | 0.32 | 0.76 | 0.76 | 0.40 |
| **MK14** | 2qd9 | 1a9u | 2.30 | 11.03 | 7.89 | 3.81 |
| **MMP13** | 830c | 1cxv | 0.30 | 1.38 | 1.38 | 1.20 |
| **MP2K1** | 3eqh | 6nyb | 0.59 | 4.11 | 3.98 | 2.62 |
| **NOS1** | 1qw6 | 1zvl | 0.16 | 6.64 | 2.00 | 1.05 |
| **NRAM** | 1b9v | 1vcj | 0.24 | 1.75 | 1.25 | 1.25 |
| **PA2GA** | 1kvo | 1kqu | 0.33 | 7.00 | 5.87 | 3.82 |
| **PARP1** | 3l3m | 2rd6 | 0.21 | 2.02 | 1.19 | 0.74 |
| **PDE5A** | 1udt | 1uho | 0.24 | 1.49 | 1.06 | 1.06 |
| **PGH1** | 2oyu | 1pgf | 0.55 | 6.31 | 5.49 | 1.75 |
| **PGH2** | 3ln1 | 1cx2 | 0.58 | 0.58 | 0.58 | 0.45 |
| **PLK1** | 2owb | 3kb7 | 0.48 | 3.17 | 3.17 | 2.17 |
| **PNPH** | 3bgs | 3phb | 0.59 | 1.12 | 1.12 | 1.12 |
| **PPARA** | 2p54 | 4ci4 | 0.58 | 1.92 | 1.54 | 1.05 |

| | | | | | | |
|---|---|---|---|---|---|---|
| **PPARD** | 2znp | 2znq | 0.33 | 2.10 | 1.74 | 1.02 |
| **PPARG** | 2gtk | 1i7i | 0.41 | 1.96 | 1.61 | 0.82 |
| **PRGR** | 3kba | 3hq5 | 0.22 | 1.10 | 1.10 | 1.10 |
| **PTN1** | 2azr | 2hb1 | 0.12 | 0.71 | 0.37 | 0.37 |
| **PUR2** | 1njs | 1rc1 | 0.11 | 1.95 | 1.30 | 1.25 |
| **PYGM** | 1c8k | 4yua | 0.18 | 4.17 | 4.17 | 2.91 |
| **PYRD** | 1d3g | 2b0m | 0.26 | 0.77 | 0.59 | 0.53 |
| **RENI** | 3g6z | 3g72 | 0.40 | 2.10 | 0.98 | 0.98 |
| **ROCK1** | 2etr | 4yvc | 0.37 | 2.05 | 1.61 | 1.34 |
| **RXRA** | 1mv9 | 4m8e | 0.21 | 1.71 | 1.68 | 1.41 |
| **SAHH** | 1li4 | 5axb | 0.15 | 0.63 | 0.63 | 0.63 |
| **SRC** | 3el8 | 3geq | 1.02 | 1.47 | 1.47 | 1.47 |
| **TGFR1** | 3hmm | 6b8y | 0.38 | 0.48 | 0.48 | 0.48 |
| **THB** | 1q4x | 1nq0 | 0.92 | 1.82 | 1.67 | 0.80 |
| **THRB** | 1ype | 1vzq | 0.19 | 0.74 | 0.65 | 0.59 |
| **TRY1** | 2ayw | 1f0u | 0.25 | 10.40 | 10.11 | 7.24 |
| **TRYB1** | 2zec | 2zeb | 0.15 | 1.13 | 0.84 | 0.60 |
| **TYSY** | 1syn | 1aiq | 0.58 | 2.42 | 2.42 | 2.18 |
| **UROK** | 1sqt | 4fu7 | 0.21 | 1.04 | 1.04 | 0.89 |
| **VGFR2** | 2p2i | 2oh4 | 2.32 | 9.88 | 9.88 | 1.14 |
| **WEE1** | 3biz | 2in6 | 0.21 | 1.97 | 1.28 | 1.10 |
| **XIAP** | 3hl5 | 1tfq | 0.79 | 3.95 | 3.90 | 3.25 |

***Table S2****: Templates used for the pharmacophore-based protocol. Column explanations are the same as for table S1.*

| TARGET | TARGET PDBID | TEMPLATE PDBID | BINDING SITE RMSD [Å] | TOP1 [Å] | TOP5 [Å] | BEST [Å] |
|---|---|---|---|---|---|---|
| **AA2AR** | 3eml | 5iua | 0.27 | 1.96 | 1.03 | 0.80 |
| **ABL1** | 2hzi | 1m52 | 0.39 | 0.70 | 0.62 | 0.62 |
| **ACE** | 3bkl | 3bkk | 0.24 | 0.74 | 0.70 | 0.54 |
| **ACES** | 1e66 | 1odc | 0.09 | 3.76 | 2.52 | 2.37 |
| **ADA** | 2e1w | 1v79 | 0.57 | 2.31 | 0.90 | 0.90 |
| **ADA17** | 2oi0 | 3b92 | 0.36 | 1.38 | 1.16 | 1.15 |
| **ADRB1** | 2vt4 | 5a8e | 0.36 | 1.02 | 0.70 | 0.41 |
| **ADRB2** | 3ny8 | 6ps5 | 0.33 | 1.55 | 1.17 | 1.00 |
| **AKT1** | 3cqw | 3mv5 | 0.34 | 0.57 | 0.57 | 0.57 |
| **AKT2** | 3d0e | 3e88 | 0.43 | 2.92 | 2.58 | 2.09 |
| **ALDR** | 2hv5 | 1iei | 0.78 | 1.80 | 0.97 | 0.89 |
| **AMPC** | 1l2s | 1xgi | 0.21 | 1.70 | 1.19 | 0.90 |
| **ANDR** | 2am9 | 1xow | 0.20 | 1.33 | 1.24 | 0.62 |
| **AOFB** | 1s3b | 2byb | 0.18 | 1.51 | 1.42 | 0.90 |
| **BACE1** | 3l5d | 3l5c | 0.11 | 5.25 | 5.21 | 4.61 |
| **BRAF** | 3d4q | 3psd | 0.34 | 2.51 | 2.02 | 1.76 |
| **CAH2** | 1bcd | 5flr | 0.11 | 2.32 | 0.76 | 0.66 |
| **CASP3** | 2cnk | 2cnl | 0.13 | 2.66 | 2.05 | 1.63 |
| **CDK2** | 1h00 | 1v1k | 0.46 | 2.00 | 1.48 | 1.31 |
| **COMT** | 3bwm | 4pyl | 0.21 | 2.76 | 2.76 | 2.60 |
| **CP2C9** | 1r9o | 5x23 | 0.74 | 0.98 | 0.69 | 0.69 |
| **CP3A4** | 3nxu | 4i4g | 0.75 | 2.05 | 2.05 | 1.50 |
| **CSF1R** | 3krj | 3dpk | 0.77 | 2.86 | 2.20 | 1.16 |
| **DEF** | 1lru | 3k6l | 0.51 | 1.49 | 0.84 | 0.84 |
| **DHI1** | 3frj | 3d4n | 0.66 | 1.84 | 1.63 | 1.42 |
| **DPP4** | 2i78 | 4j3j | 0.25 | 1.56 | 1.14 | 0.72 |
| **DYR** | 3nxo | 3gyf | 0.33 | 1.55 | 0.62 | 0.58 |
| **EGFR** | 2rgp | 3bel | 0.11 | 1.00 | 1.00 | 0.85 |
| **ESR1** | 1sj0 | 1yim | 0.43 | 0.89 | 0.77 | 0.66 |
| **ESR2** | 2fsz | 1l2j | 0.82 | 3.82 | 3.45 | 2.37 |
| **FA10** | 3kl6 | 2cji | 0.34 | 1.47 | 1.42 | 1.31 |
| **FA7** | 1w7x | 4jyu | 2.26 | 4.17 | 1.46 | 1.13 |
| **FABP4** | 2nnq | 5d47 | 0.37 | 5.95 | 5.45 | 3.21 |
| **FAK1** | 3bz3 | 4d58 | 1.39 | 1.96 | 1.87 | 1.51 |
| **FGFR1** | 3c4f | 4wun | 0.67 | 1.10 | 0.88 | 0.81 |
| **FKB1A** | 1j4h | 1j4i | 0.44 | 2.57 | 2.57 | 2.20 |
| **FNTA** | 3e37 | 3e33 | 0.21 | 5.67 | 3.96 | 2.43 |
| **FPPS** | 1zw5 | 2opm | 0.30 | 1.68 | 1.67 | 1.18 |
| **GCR** | 3bqd | 3mnp | 0.51 | inf | 0.54 | 0.54 |
| **GLCM** | 2v3f | 3rik | 1.48 | 3.38 | 2.03 | 1.70 |
| **GRIA2** | 3kgc | 3ilu | 2.03 | 17.16 | 15.86 | 15.39 |
| **GRIK1** | 1vso | 2qs1 | 0.97 | 2.45 | 2.17 | 1.72 |
| **HDAC2** | 3max | 4ly1 | 0.11 | 1.65 | 1.40 | 1.04 |

| | | | | | | |
|---|---|---|---|---|---|---|
| **HDAC8** | 3f07 | 1t64 | 1.35 | 4.65 | 4.65 | 3.77 |
| **HIVINT** | 3nf7 | 3nf9 | 0.17 | 2.87 | 2.87 | 2.75 |
| **HIVPR** | 1xl2 | 1xl5 | 1.40 | 6.55 | 6.33 | 5.55 |
| **HIVRT** | 3lan | 3lam | 0.38 | 2.29 | 2.26 | 0.58 |
| **HMDH** | 3ccw | 3ccz | 0.14 | 0.65 | 0.65 | 0.65 |
| **HS90A** | 1uyg | 1uyc | 0.16 | 2.30 | 2.15 | 1.88 |
| **HXK4** | 3f9m | 3goi | 0.87 | 0.95 | 0.50 | 0.50 |
| **IGF1R** | 2oj9 | 2zm3 | 0.86 | 7.64 | 3.10 | 2.58 |
| **INHA** | 2h7l | 4tzk | 0.12 | 1.01 | 0.64 | 0.64 |
| **ITAL** | 2ica | 3m6f | 0.27 | 2.13 | 1.68 | 0.99 |
| **JAK2** | 3lpb | 3krr | 0.40 | 1.60 | 1.46 | 0.57 |
| **KIF11** | 3cjo | 2fky | 0.31 | 1.86 | 1.86 | 1.57 |
| **KIT** | 3g0e | 6mob | 0.20 | 10.61 | 6.40 | 3.94 |
| **KITH** | 2b8t | 2uz3 | 0.46 | 1.65 | 0.61 | 0.60 |
| **LCK** | 2of2 | 2of4 | 0.11 | 1.24 | 0.67 | 0.64 |
| **LKHA4** | 3chp | 3cho | 0.16 | 2.06 | 1.78 | 0.94 |
| **MAPK2** | 3m2w | 3m42 | 0.66 | 1.03 | 0.81 | 0.76 |
| **MCR** | 2aa2 | 2a3i | 0.18 | 0.61 | 0.60 | 0.43 |
| **MET** | 3lq8 | 5hti | 0.22 | 0.89 | 0.84 | 0.80 |
| **MK01** | 2ojg | 2oji | 0.95 | 1.90 | 1.36 | 1.30 |
| **MK10** | 2zdt | 4g1w | 0.78 | 0.98 | 0.98 | 0.63 |
| **MK14** | 2qd9 | 3ha8 | 1.38 | 2.14 | 1.89 | 1.08 |
| **MMP13** | 830c | 2pjt | 0.27 | 1.80 | 1.48 | 1.17 |
| **MP2K1** | 3eqh | 3zlx | 0.66 | 11.41 | 11.07 | 10.06 |
| **NOS1** | 1qw6 | 3hsn | 0.16 | 2.28 | 1.80 | 1.16 |
| **NRAM** | 1b9v | 1vcj | 0.24 | 1.41 | 1.12 | 1.12 |
| **PA2GA** | 1kvo | 1kqu | 0.33 | 3.06 | 1.97 | 1.78 |
| **PARP1** | 3l3m | 6vko | 0.61 | 1.36 | 0.37 | 0.37 |
| **PDE5A** | 1udt | 1xp0 | 2.86 | 3.20 | 2.90 | 2.61 |
| **PGH1** | 2oyu | 1ht5 | 0.37 | 7.28 | 6.82 | 6.21 |
| **PGH2** | 3ln1 | 6bl4 | 0.36 | 1.55 | 1.40 | 1.26 |
| **PLK1** | 2owb | 3kb7 | 0.48 | 5.60 | 5.56 | 2.42 |
| **PNPH** | 3bgs | 4ear | 0.62 | 1.35 | 1.16 | 1.04 |
| **PPARA** | 2p54 | 1kkq | 4.01 | 8.43 | 8.27 | 6.96 |
| **PPARD** | 2znp | 2znq | 0.33 | 0.86 | 0.86 | 0.86 |
| **PPARG** | 2gtk | 3ia6 | 0.46 | 1.57 | 0.77 | 0.77 |
| **PRGR** | 3kba | 2ovh | 1.20 | 6.22 | 5.02 | 4.38 |
| **PTN1** | 2azr | 4yua | 0.18 | 0.70 | 0.49 | 0.49 |
| **PUR2** | 1njs | 1uuo | 0.82 | 2.34 | 2.33 | 1.16 |
| **PYGM** | 1c8k | 3g72 | 0.39 | 5.53 | 5.45 | 3.08 |
| **PYRD** | 1d3g | 5wng | 0.53 | 0.70 | 0.70 | 0.50 |
| **RENI** | 3g6z | 4rmd | 0.20 | 1.10 | 0.75 | 0.75 |
| **ROCK1** | 2etr | 5axc | 0.11 | 9.62 | 9.50 | 7.75 |
| **RXRA** | 1mv9 | 3el7 | 1.30 | 2.72 | 2.72 | 1.73 |
| **SAHH** | 1li4 | 6b8y | 0.38 | 0.71 | 0.69 | 0.61 |
| **SRC** | 3el8 | 3imy | 0.66 | 1.07 | 0.94 | 0.56 |
| **TGFR1** | 3hmm | 1ypg | 0.15 | 0.64 | 0.64 | 0.64 |
| **THB** | 1q4x | 1f0u | 0.25 | 1.97 | 1.97 | 0.88 |

| | | | | | | |
|---|---|---|---|---|---|---|
| **THRB** | 1ype | 2zeb | 0.15 | 0.71 | 0.61 | 0.56 |
| **TRY1** | 2ayw | 2kce | 0.55 | 11.23 | 6.39 | 5.66 |
| **TRYB1** | 2zec | 4fu7 | 0.21 | 2.16 | 2.16 | 1.41 |
| **TYSY** | 1syn | 3efl | 0.27 | 3.18 | 2.29 | 1.70 |
| **UROK** | 1sqt | 3bi6 | 0.13 | 2.21 | 1.72 | 1.22 |
| **VGFR2** | 2p2i | 1tft | 0.61 | 1.23 | 1.23 | 1.01 |
| **WEE1** | 3biz | 1rc1 | 0.11 | 1.32 | 1.23 | 0.84 |
| **XIAP** | 3hl5 | 2hb1 | 0.12 | 4.63 | 3.38 | 3.38 |

*Table S3: HADDOCK run.cns parameters that were modified for these protocols.*

| **Setting** | **Value** |
|---|---|
| ncomponents | 3 |
| fix_origin_mol1[1] | true |
| fix_origin_mol3[2] | true |
| shape_mol3 | true |
| delenph | false |
| amb_lastit | 1 |
| noecv | false |
| prot_top_mol3 | shape.top |
| prot_par_mol3 | shape.param |
| dielec_0 | cdie |
| epsilon_1 | 10 |
| dielec_1 | cdie |
| inter_rigid | 0.001 |
| rotate180_it0 | false |
| firstwater | no |
| w_vdw_0 | 0 |
| w_elec_2 | 0.1 |
| w_dist_2 | 0 |
| clust_meth | RMSD |
| clust_cutoff | 1.5 |

[1] Refers to the receptor molecule.
[2] Refers to the shape beads.

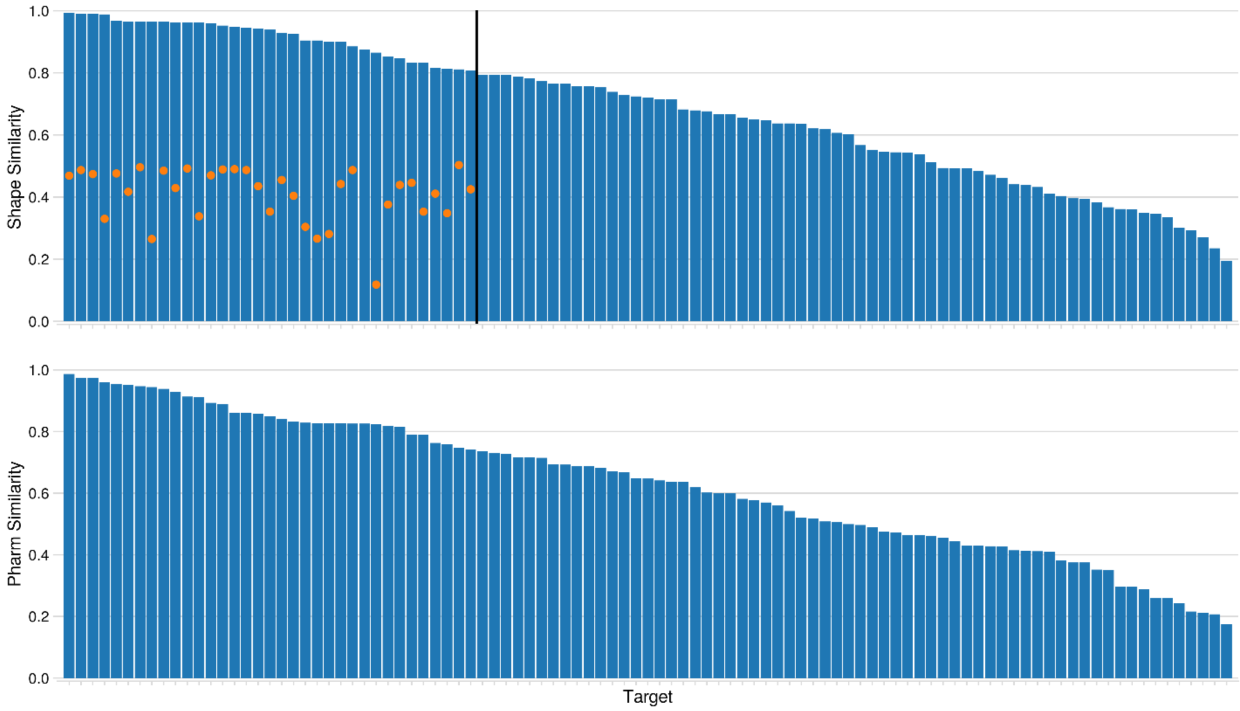


***Figure S1***: *Distribution of template compound similarity values for shape (top) and pharm templates (bottom). The similarity metric used for the shape-based protocol is the Tversky coefficient computed over the Maximum Common Substructure and for the pharmacophore-based protocol is the Tanimoto coefficient computed over 2D pharmacophore fingerprints. The height of the blue bar indicates the similarity value for a given target. The orange dots of the shape chart indicate the similarity of the low similarity compound that was chosen for that particular target. Black line in the top plot indicates the targets for which the identified templates have a similarity of more than 0.8 to their respective reference. Note: The two plots have been sorted according to their respective similarity metric, meaning the bars do not (necessarily) correspond to the same target between the top and bottom plots.*

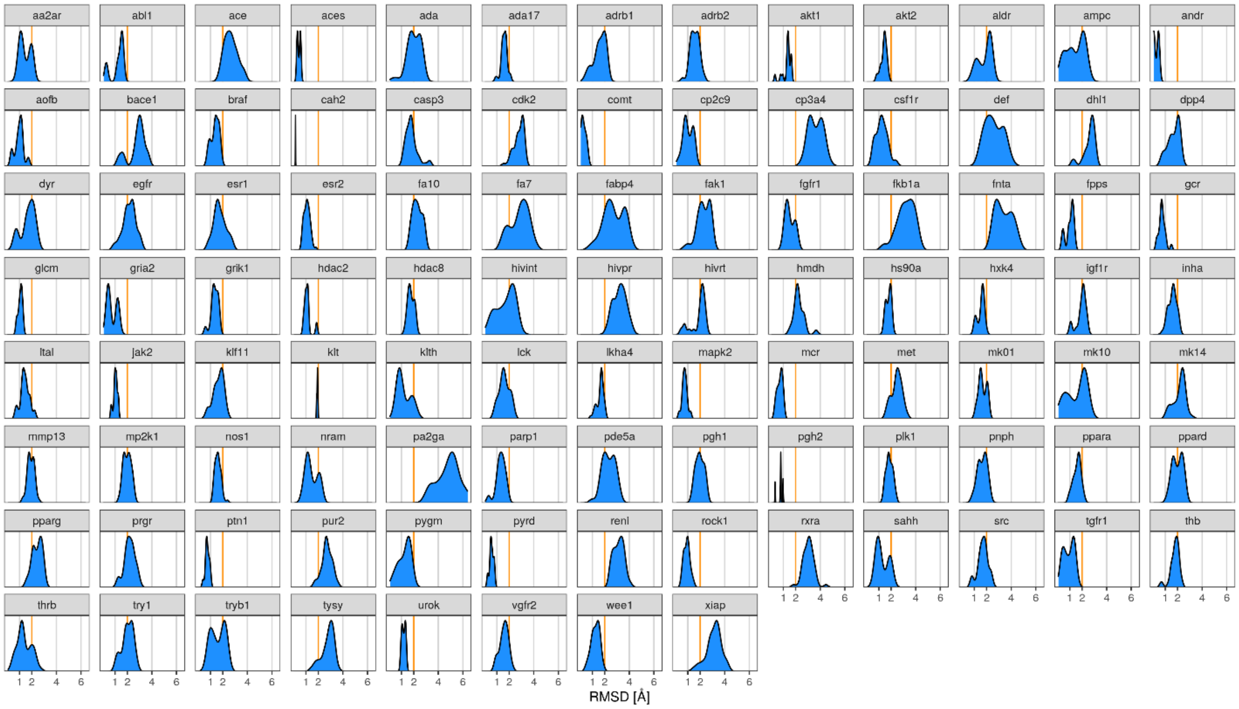


***Figure S2**: Distributions of RMSD values of the conformers generated with RDKit relative to the reference compound.*

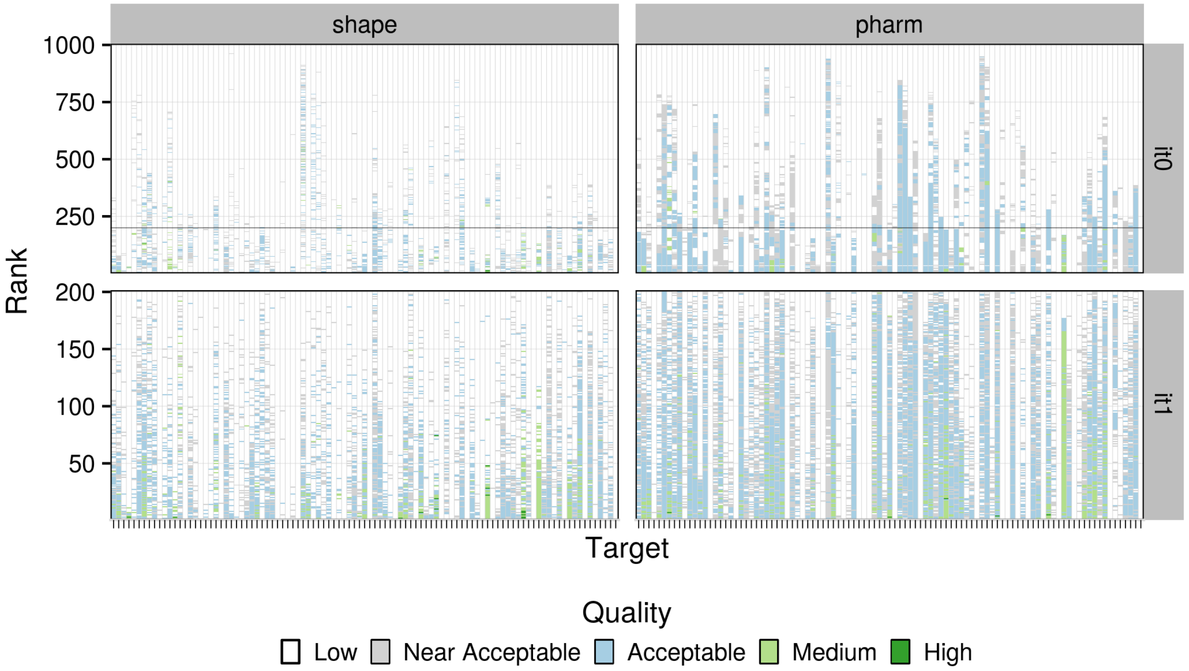


***Figure S3***: *Assessment of the performance of the two protocols for the two modelling stages. Results of the rigid-body docking (it0; top panel) and semi-flexible refinement (it1; bottom panel) are shown. The Y axis for all sub-graphs corresponds to the ranking of the models according to the HADDOCK scoring function, with models ranked near 0 having the best scores. Every model has been coloured according to its quality, with high-, medium-, acceptable-, near acceptable- and low-quality models having IL-RMSD values of less than 0.5 Å (dark green), between 0.5 and 1 Å (light green), between 1 and 2 Å (light blue), between 2 and 2.5 Å (light grey) and over 2.5 Å (white), respectively. The black line indicates the threshold for it0 models to proceed to refinement.*

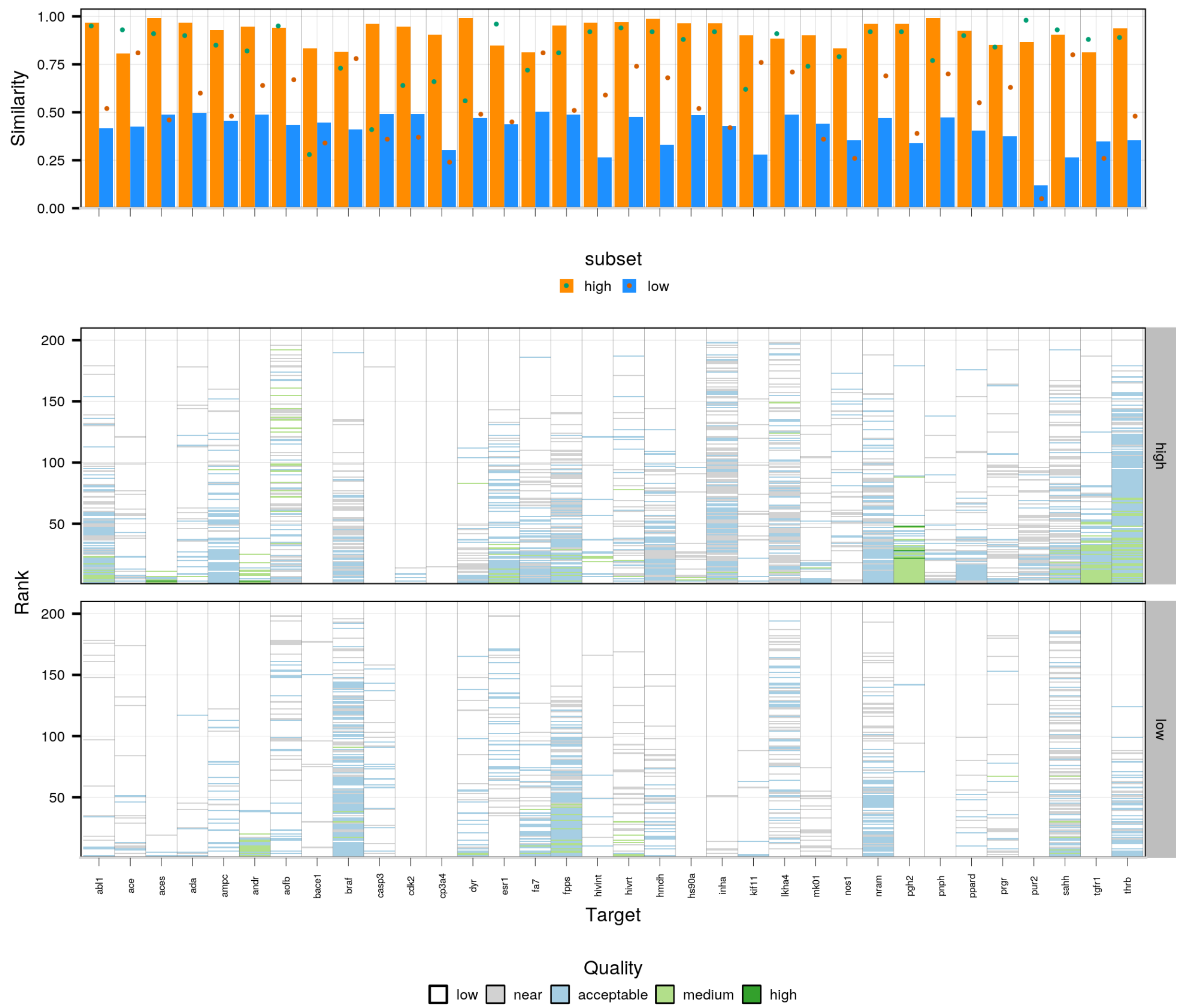


***Figure S4***: *Comparison of template quality (Tversky similarity computed over Maximum Common Substructure) and performance for the low- and high-quality subsets for the shape-based protocol. The top panel highlights the similarity (orange and blue bars for the high- and low-similarity subsets, respectively) and overlap (green and dark green dots for the high- and low-similarity subsets, respectively) between reference and template compounds. The bottom panel compares the performance for the semi-flexible refinement stage of HADDOCK (it1) for the two subsets. Each column corresponds to one target with the Y axis reflecting the ranking of models (ranks close to 0 refer to top-ranked models and those close to 200 to bottom-ranked models) and the colour of each model reflecting its quality (see description of figure S3).*

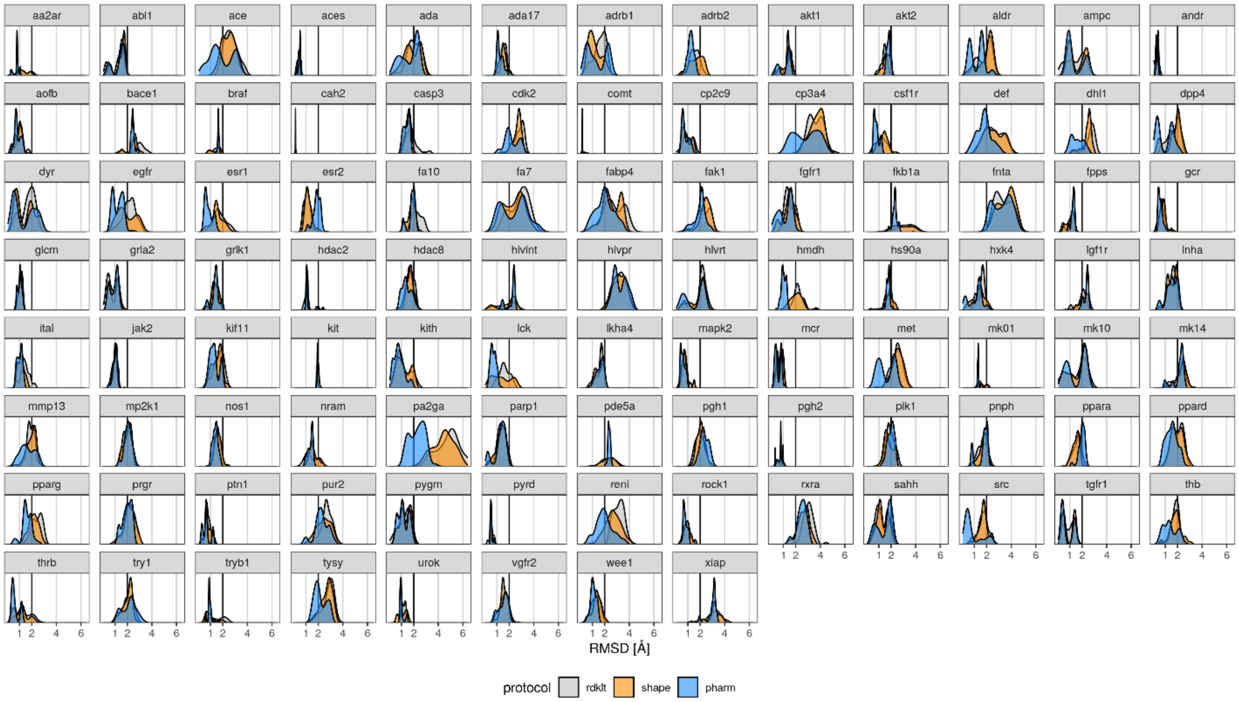


***Figure S5***: *Distributions of model compound RMSD values compared to their respective reference compounds. Distributions shaded in light grey, orange and blue correspond to the compounds generated by RDKit, and the compounds at the end of the refinement stage of the shape-based and pharmacophore-based protocols, respectively.*

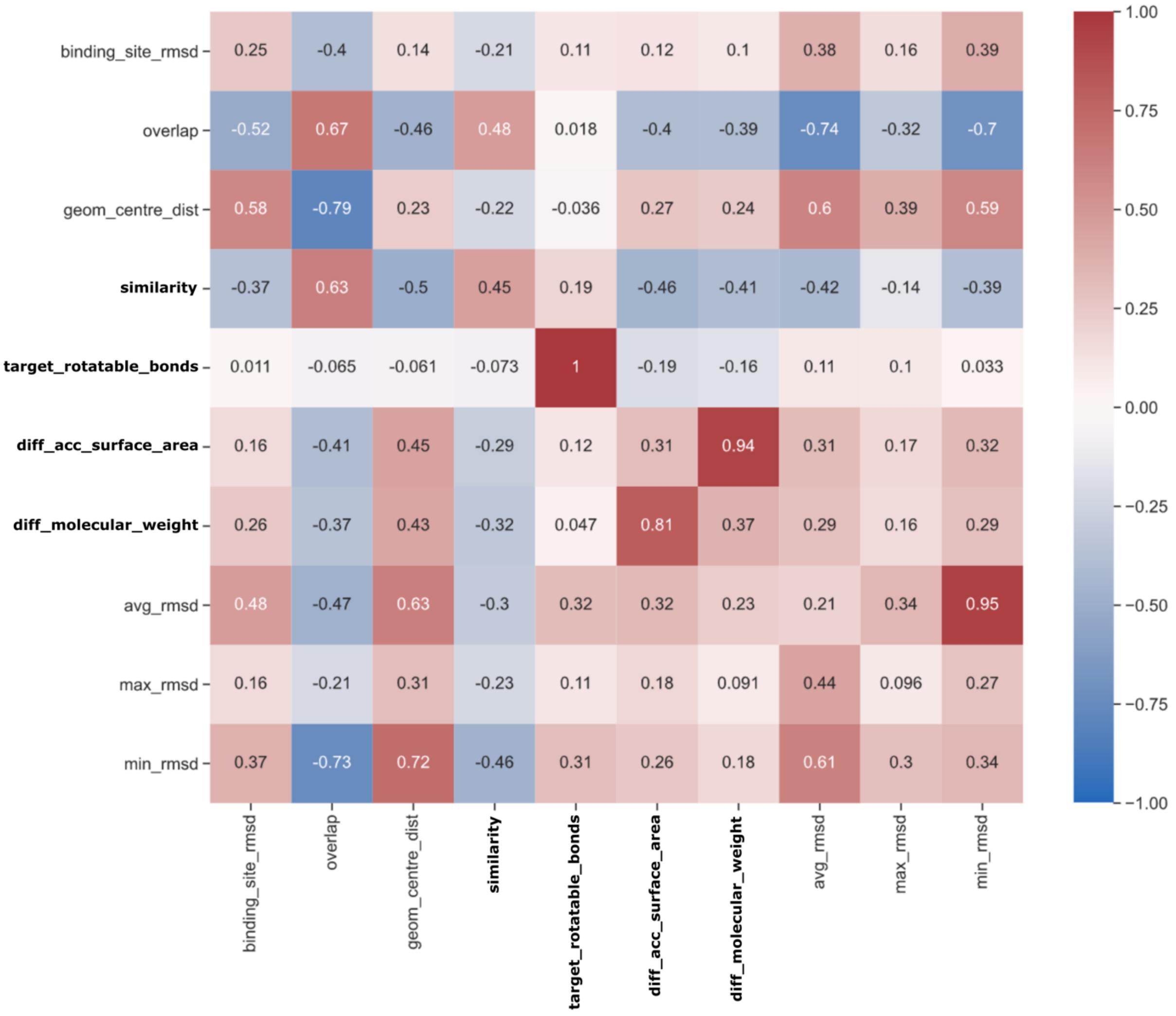


***Figure S6***: *Heatmap of the correlation between different metrics computed on the template and target compounds used for each protocol and the* ***it1*** *results. The lower part of the matrix (below the diagonal) contains correlation information between the metrics of the shape-based protocol and the upper part the correlations for the pharmacophore-based protocol. For example, the last column of the heatmap contains all correlation values between the minimum RMSD per target obtained and all the other metrics for the pharmacophore-based protocol, and the last row all correlation values between the minimum RMSD per target obtained and all the other metrics for the shape-based protocol. The diagonal represents correlation between the shape-based protocol and the pharmacophore protocol metrics.*

*The binding_site_rmsd is computed as the RMSD between all residues within 5A of the bound compound in the target and the template receptor structures; The overlap is computed with the Exact Overlap metric of the shape toolkit of OpenEye (release 2020.2.0) after superimposing on the backbone atoms of the binding site of the receptors; the geom_centre_dist is the distance between the template compound and the target compound geometrical centre once the receptors are superimposed; the similarity stands for Tanimoto coefficient in the vertical axis, and Tversky similarity in the horizontal axis; the number of rotatable bond, the Labute accessible surface area and the difference in molecular weight were computed with RDKIT; The avg_rmsd, min_rmsd and max_rmsd stand for the average, minimum and maximum IL-RMSD obtained per target at it1. The metrics in bold are predictive metrics which can be calculated without the knowledge of the reference complex.*

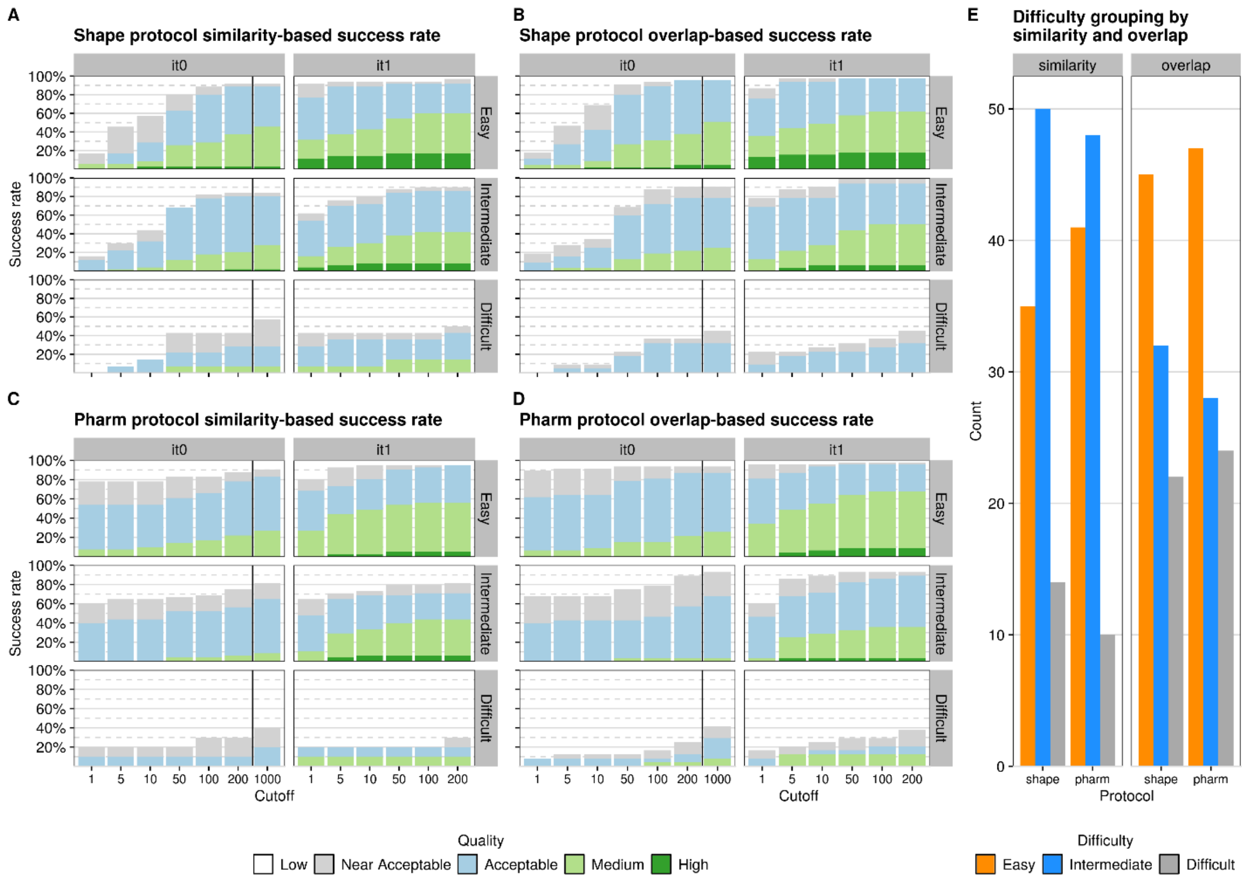


***Figure S7**: Evaluation of the success rate of the two protocols (shape-based: panels A & B and pharmacophore-based: panels C & D) as a function of compound similarity (panels A & C) and overlap (panels B & D) and breakdown of the targets in difficulty tiers according to similarity and overlap (panel E). Targets with Tv greater than 0.8, between 0.4 and 0.8, and below 0.4 are classified as "Easy", "Intermediate" and "Difficult", respectively, for the shape-based protocol. Targets with Tc greater than 0.7, between 0.3 and 0.7, and below 0.3 are classified as "Easy", "Intermediate" and "Difficult", respectively, for the pharmacophore-based protocol. Targets with overlap greater than 0.75, between 0.5 and 0.75, and below 0.5 are classified as "Easy", "Intermediate" and "Difficult", respectively, for both protocols.*

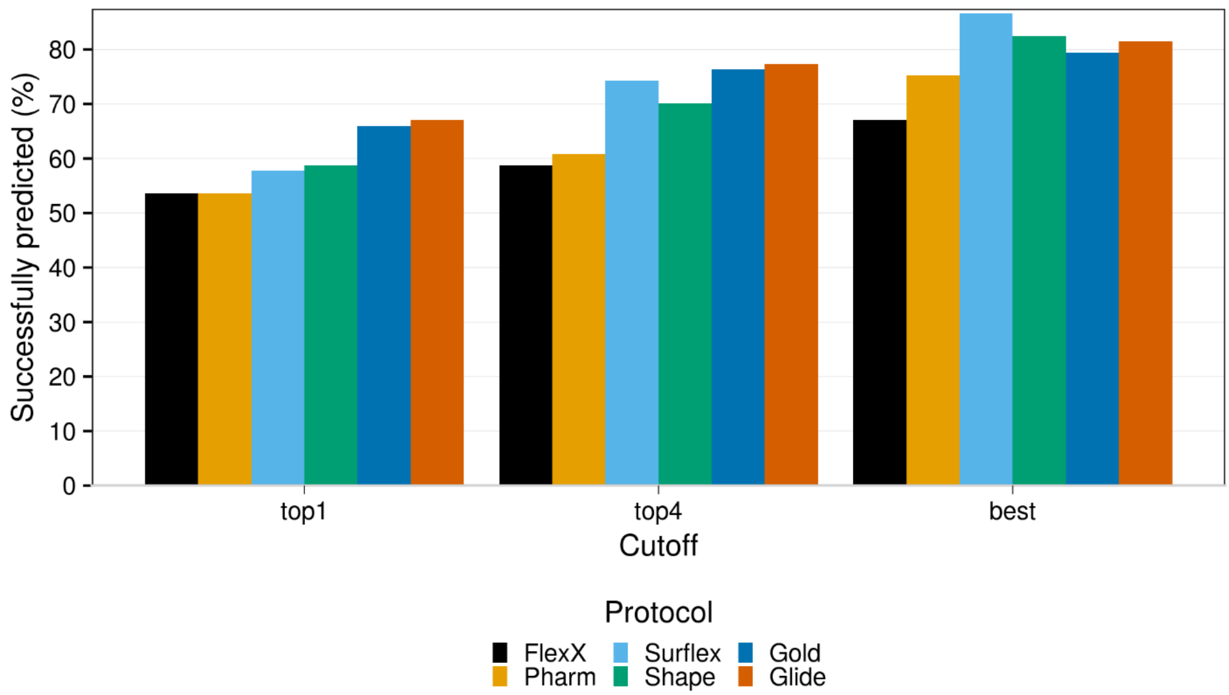


***Figure S8**: Success rate of various commercial docking platforms against our two protocols evaluated as a function of the top-ranked models, the top4 ranked models and all models generated.*
